# Microbiota-root-shoot axis modulation by MYC2 favours *Arabidopsis* growth over defence under suboptimal light

**DOI:** 10.1101/2020.11.06.371146

**Authors:** Shiji Hou, Thorsten Thiergart, Nathan Vannier, Fantin Mesny, Jörg Ziegler, Brigitte Pickel, Stéphane Hacquard

## Abstract

Bidirectional root-shoot signalling is likely key in orchestrating stress responses and ensuring plant survival. Here we show that *Arabidopsis thaliana* responses to microbial root commensals and light are interconnected along a microbiota-root-shoot axis. Microbiota and light manipulation experiments in a gnotobiotic system reveal that low photosynthetically active radiation (LP) perceived by leaves induce long-distance modulation of root bacterial, but not fungal or oomycetal communities. Reciprocally, bacterial root commensals and particularly *Pseudomomas* isolates are necessary for rescuing plant growth under LP. RNA-Seq, combined with leaf inoculation experiments with biotrophic and necrotrophic microbial pathogens indicate that microbiota-induced growth under LP coincides with transcriptional repression of immune responses, thereby increasing susceptibility to both pathogens. Inspection of a set of *A. thaliana* mutants demonstrates that orchestration of this light-dependent growth-defence trade-off requires the transcriptional regulator MYC2. Our work indicates that aboveground stress responses in plants can be governed by signals from microbial root commensals.

## INTRODUCTION

Unlike animals, plants are sessile organisms that must simultaneously integrate various responses to biotic and abiotic stresses to prioritize either growth or defence depending on rapidly changing surrounding conditions (**Huot et al., 2014**). Since colonization of land by ancestral plant lineages 450 MyA, complex multi-kingdom microbial consortia interact with and colonize roots of healthy plants (**Hassani et al., 2018**). Therefore, direct integration of microbial and environmental cues by plants is likely key in orchestrating their ability to overcome environmental stresses (**Hacquard et al., 2016; Castrillo et al., 2017; Stringlis et al., 2018**). The beneficial activities conferred by microbial root commensals have been primarily described in the context of nutritional stress (**Hiruma et al., 2016; Castrillo et al., 2017; Almario et al., 2017; Zhang et al., 2019; Harbort et al., 2020**) or pathogen attack (**Kwak et al., 2018; Berendsen et al., 2018; Durán et al., 2018; Carrión et al., 2019; Gu et al., 2020**), revealing that root colonization by complex microbial communities can shape host phenotypic plasticity and protect plants from diseases.

Carbohydrate biosynthesis, energy production, and therefore plant growth depends on the amount and quality of photosynthetically active radiation (PAR) absorbed by chloroplasts in leaves (**Poorter et al., 2019**). Spectral PAR irradiance ranges from 400 to 700 nm under natural conditions, and a reduction of both the amount of PAR (light quantity) and the ratio of red to far-red light (light quality) is perceived as a warning signal in plants, thereby triggering adaptive changes in shoot morphology (**Franklin and Whitelam 2005; Franklin 2008**). Light-dependent resource allocation was shown to be tightly linked with defence signalling in *A. thaliana* leaves, suggesting that aboveground responses to light and pathogens are interconnected (**Griebel and Zeier 2008; Leone et al., 2014; Campos et al., 2016; Fernández-Milmanda et al., 2020; Courbier et al., 2020**). However, the extent to which belowground commensals can modulate, or even dictate aboveground stress responses remains unknown. Given the fact that a substantial amount of photosynthetically fixed carbohydrates (i.e., > 15%) is invested in the rhizosphere (**Haichar et al., 2008; Hernández et al., 2015**), we hypothesized that the aboveground response to light and the belowground response to microbes are interconnected.

Here, we show that microbial root commensals can alleviate *A. thaliana* growth deficiency under suboptimal light and we demonstrate that this growth rescue is explained by a prioritization of microbiota-induced growth over defence responses in shoot. Our results indicate that the two major functions conferred by root commensals (shoot growth promotion and shoot defence priming) are orchestrated through the transcriptional regulator MYC2 to either prioritize growth or defence depending on the surrounding light conditions. This implies that the belowground response to microbes and the aboveground response to light are integrated along a microbiota-root-shoot axis, thereby promoting the plant’s ability to alleviate an abiotic stress.

## RESULTS

### A synthetic root microbiota rescued Arabidopsis growth under low PAR

We hypothesized that a microbiota-root-shoot axis exists in plants, allowing them to take advantage of belowground microbial commensals to orchestrate aboveground stress responses. First, we assembled a synthetic microbial consortium (SynCom, **Figure S1A, Table S1**) with phylogenetically diverse bacteria (B, 183 strains), fungi (F, 24 strains), and oomycetes (O, 7 strains) that naturally colonize roots of healthy *A. thaliana* (**Durán et al., 2018; Thiergart et al., 2020**). Eight versions of this BFO SynCom containing B, F, and O communities mixed at different relative abundance (RA) in the starting inoculum were used to re-colonize roots of germ-free *A. thaliana* in the gnotobiotic FlowPot system (**Kremer et al., 2016**). Sequencing of the 16S rRNA gene (B) and ITS (F, O) confirmed that the initial difference in RA between the three microbial groups did not affect output root microbiota assembly five weeks post-inoculation, based on principal component analysis (**Figure S1B**) and constrained analysis of principal coordinates (CAP, B: *P* = 0.921, F: *P* = 0.476, O: *P* = 0.075). Similarly, difference in the B-to-F-to-O RA did not modulate the BFO-mediated effect on plant growth (Kruskal-Wallis, *P* = 0.226, **Figure S1C**). We then tested the relevance of this multi-kingdom microbial SynCom for plant growth under suboptimal light conditions. We manipulated light conditions in our plant growth chamber 1) by restricting shoot exposure to direct light (Low PAR, LP, **Figures S1D** and **S1E**), resulting in ~55% reduction in photosynthetic photon flux density (PPFD 400-700nm, NC = 62.35, LP = 27.91, Mann-Whitney U test, *P*< 2.2e-16, **Figure S1D**) while only marginally affecting the R:FR ratio (NC = 9.24, LP = 9.14, **Figure S1F**) and the temperature (NC = 21.02°C, LP = 20.90°C, **Figure S1F**) and 2) by exposing shoot to far red light (740 nm) for 15 minutes at the end of the day (EODFR, **Dubois et al., 2010**). Compared to five-week-old plants grown under normal light conditions (NC), those exposed to LP and EODFR showed a significant reduction in canopy size (LP: 2.4-fold, *P* = 5.45e-13; EODFR: 2.1-fold, *P* = 2.88e-4; Mann-Whitney-U test, **Figures 1A, 1B** and **S1G**) and shoot fresh weight (LP: 2.2-fold, *P* = 2.32e-8; EODFR: 2.2-fold, *P* = 4.11e-5; Mann-Whitney-U test, **Figure 1C**) in the absence of the BFO SynCom. Remarkably, the presence of BFO rescued LP- and EODFR-mediated decrease in canopy size and shoot fresh weight to control levels (**Figures 1A-C**). Furthermore, recolonization experiments with B in the presence or absence of F and O communities indicated that B commensals alone partially rescued plant growth under LP, whereas the synergistic contribution of B and filamentous eukaryotes was needed for the full rescue (Kruskal-Wallis with Dunn’s *post hoc* test, *P* < 0.05, **Figure 1D**). The presence of BFO also led to a significant increase in petiole length (**Figure 1E**), leaf number (**Figure 1F**), and the leaf length/width ratio (i.e. a proxy of leaf shape, **Figure 1G**) (Kruskal-Wallis with Dunn’s *post hoc* test, *P* < 0.05). Inspection of the magnitude of BFO-mediated modification of leaf traits indicated quantitative differences among the tested parameters and revealed that the amplitude of the BFO effect was always greater under LP and EODFR than under the control light condition (Cohen’s effect size, **Figure 1H**). Our results indicate that belowground microbial commensals promote the plant’s ability to grow under suboptimal light conditions by promoting canopy size and to a lesser extent by modulating canopy shape.

**Figure 1.**
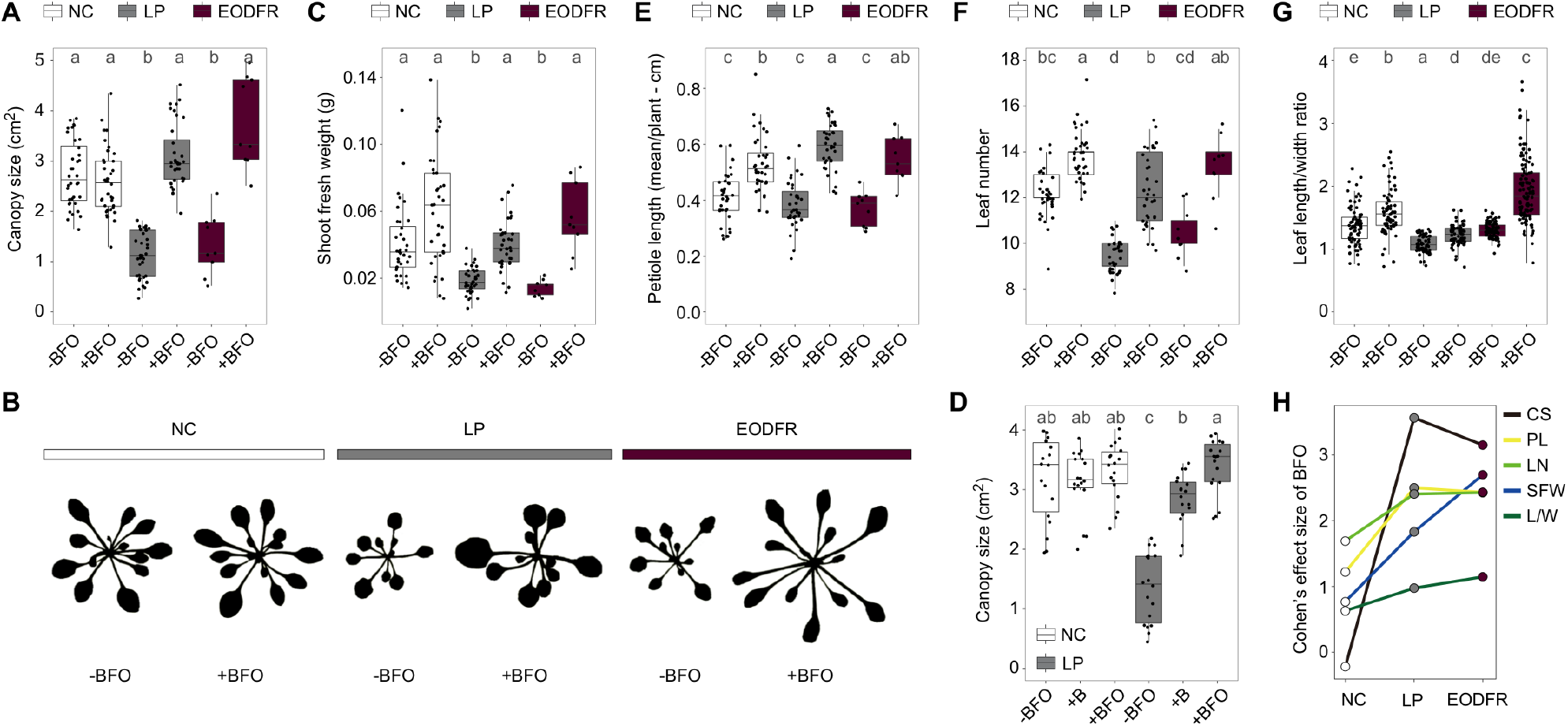
BFO-mediated modulation of leaf traits under LP and EODFR. **(A)** Canopy size (in cm^2^) of five-week-old *A. thaliana* grown in the FlowPot system in the absence (-) or presence (+) of a synthetic microbial community (BFO SynCom, B = 183 bacteria, F = 24 fungi, O = 7 oomycetes) under normal light conditions (NC, white), low photosynthetically active radiation (LP, grey), or end-of-day far-red (EODFR, dark red). Three independent biological replicates (n = 162 plants). Letters indicate statistical significance corresponding to Kruskal-Wallis with Dunn’s *post hoc* test (α = 0.05). **(B)** Representative images illustrating light- and BFO-induced change in shoot morphology. **(C)** Shoot fresh weight (in g) of five-week-old *A. thaliana* inoculated with and without the BFO SynCom under NC, LP, and EODFR. Three independent biological replicates (n = 161 plants). Letters indicate statistical significance corresponding to Kruskal-Wallis with Dunn’s *post hoc* test (α = 0.05). **(D)** Canopy size (in cm^2^) of five-week-old *A. thaliana* grown in the FlowPot system in the absence (-) or presence (+) of a synthetic microbial community composed of either B or BFO under NC (white) and LP (grey). Three independent biological replicates (n = 108 plants). Letters indicate statistical significance corresponding to Kruskal-Wallis with Dunn’s *post hoc* test (α = 0.05). **(E)** Mean petiole length (in cm) of five-week-old *A. thaliana* inoculated with and without the BFO SynCom under NC, LP, and EODFR. Three independent biological replicates (n = 161 plants). Letters indicate statistical significance corresponding to ANOVA followed by *post hoc* Tukey’s HSD (α = 0.05). **(F)** Total leaf number of five-week-old *A. thaliana* inoculated with and without the BFO SynCom under NC, LP, and EODFR. Three independent biological replicates (n = 161 plants). Letters indicate statistical significance corresponding to Kruskal-Wallis with Dunn’s *post hoc* test (α = 0.05). **(G)** Leaf length/width ratio of five-week-old *A. thaliana* inoculated with and without the BFO SynCom under NC, LP, and EODFR. Ratios close to one reflect round leaf shape, whereas ratios >1 reflect elongated leaf shape. Three independent biological replicates (n = 486 leaves). Letters indicate statistical significance corresponding to Kruskal-Wallis with Dunn’s *post hoc* test (α = 0.05). **(H)** Effect size of BFO on shoot morphological traits under NC, LP, and EODFR. Effect size was computed between-BFO and +BFO conditions for each light condition by adjusting the calculation of the standard deviation with weights of the sample sizes (Cohen’s d). Different line colors reflect different shoot morphological traits. CS: canopy size, PL: petiole length (mean/plant), LN: leaf number, SFW: shoot fresh weight, L/W: leaf length/width ratio.

### Aboveground light conditions modulate root microbiota assembly

We tested whether changes in light conditions perceived by leaves can cascade along the shoot-root axis, thereby modulating root microbiota assembly. Using the same strategy described above, we monitored B, F, and O community composition in roots and surrounding peat matrix for both LP and NC using amplicon sequencing (see methods). The 183 B, 24 F and 7 O, respectively, could be discriminated into 115, 24, and 7 strain variants at single nucleotide resolution against reference 16S rRNA and ITS sequences (**Table S1**). Inspection of B, F, and O strain variants detected across conditions indicated a weak or not significant effect of the light condition on microbial alpha diversity (ANOVA followed by *post hoc* Tukey’s HSD, **Figure 2A**). Principal coordinates analysis of Bray-Curtis dissimilarities (PCoA, **Figure S2A**), combined with permutational multivariate analysis of variance (PERMANOVA, **Table S2**), revealed that the factor “compartment” explained a significant part of the variance in B and F, but not O community composition in our gnotobiotic system (“compartment”, B: R^2^ = 0.40, *P* < 0.001, F: R^2^ = 0.13, *P* < 0.001; O: R^2^ = 0.02, *P* = 0.439, **Table S2**). Overall, the factor “light” did not affect F and O community composition but did significantly shape B community composition (“light”, R^2^ = 0.07, *P* = 0.002, **Table S2**). CAP analysis constrained by “light” and conducted independently for root and matrix samples indicated a significant effect of the light condition on B community composition in root (“light”, 11.9 %, *P* < 0.001), but not surrounding peat samples (“light”, *P* = 0.606, **Figure 2B**). In contrast, “light” had no significant effect on F and O community composition in roots (“light”, F: *P* = 0.105, O: *P* = 0.574) and a weak impact on F community in the matrix (“light”, F: *P* = 0.023, O: *P* = 0.659, **Figure 2B**). Partial least squares-discriminant analysis (PLS-DA, **Figure 2C**), together with PERMANOVA conducted on root and matrix samples separately (**Table S2**) further validated the prominent effect of the light condition on the composition of root-associated B (“light”, root: R^2^ = 0.23, *P* = 0.006; matrix R^2^ = 0.05, *P* = 0.3), but not F and O communities (**Table S2**). Pairwise-enrichment tests conducted between LP and NC conditions revealed a significant impact of the light condition on the RA of 23 strain variants in root samples and four in matrix samples (edgeR, generalized linear model, *P* < 0.05, **Figures 2D** and **2E**). Notably, multiple *Pseudomonas* isolates were specifically enriched in the root compartment between LP and NC (**Figures 2E** and **S2B**). Given the possible ectopic leaf colonization by root microbiota members in our gnotobiotic system (**Bai et al., 2015**), we also tested whether BFO root commensals can be detected in leaves and the extent to which their RA can be modulated by light conditions. Diversity analysis (**Figure S2C**), PCoA (**Figure S2D**) and enrichment tests (**Figures S2E** and **S2F**) revealed that 44%, 15%, and 0% of B, F and O strain variants detected belowground were colonizing aboveground leaf tissues, and that the composition of ectopic B assemblage was also modulated by light conditions, with 13 strain variants showing light-dependent differential growth (**Figure S2F**). Our results indicate that shoot exposure to suboptimal light triggers host-dependent modulation of the growth of root- and ectopic leaf-associated bacterial taxa in a complex multikingdom synthetic microbiome.

**Figure 2.**
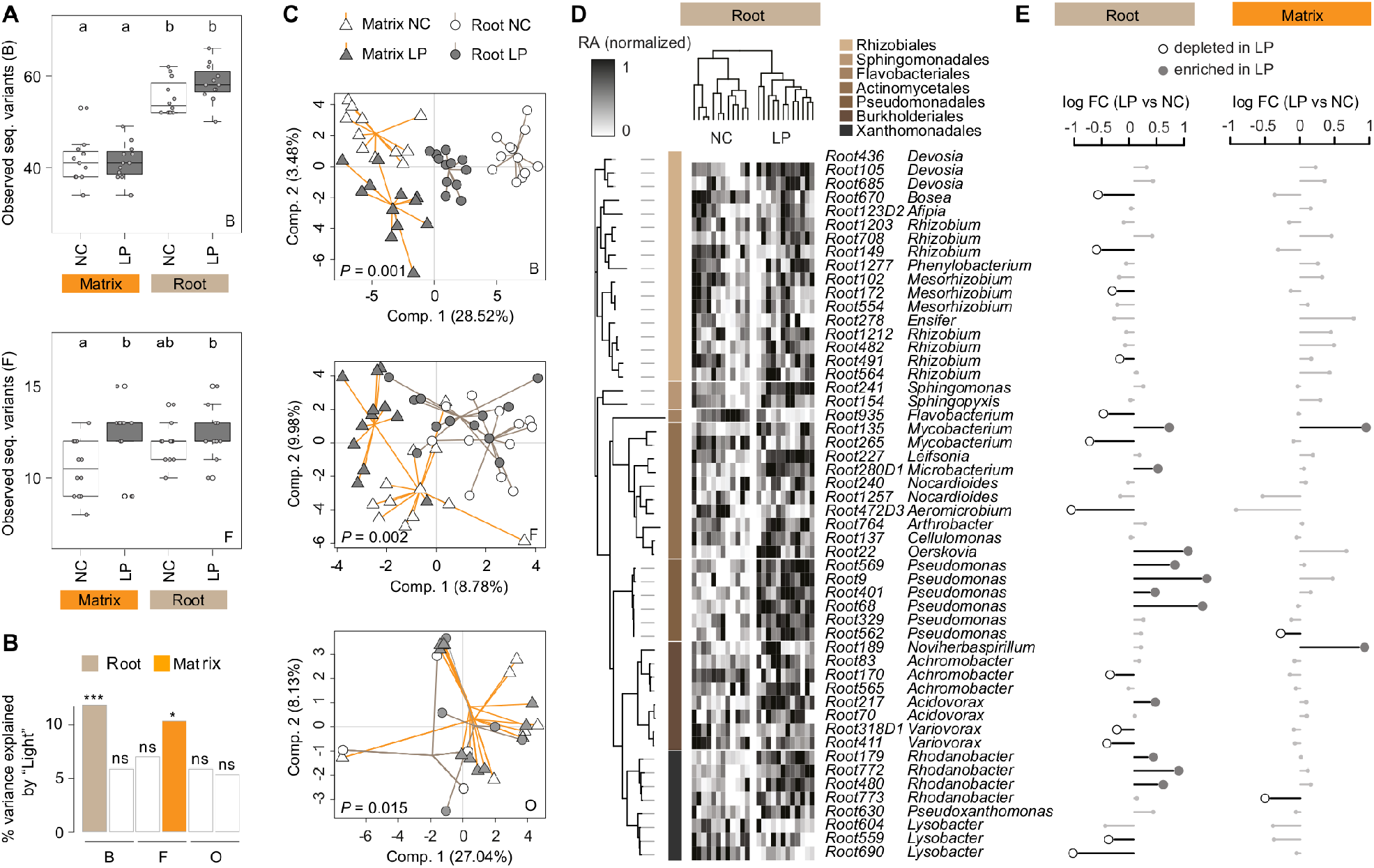
LP-mediated modulation of root microbiota assembly. **(A)** Number of bacterial (B, top) and fungal (F, bottom) strain variants detected in root and peat matrix five weeks post BFO inoculation in the FlowPot system. The 183 B, 24 F and 7 oomycetal (O) strains were grouped into 115 B, 24 F, and 7 O strain variants that could be discriminated at single nucleotide resolution against reference 16S rRNA and ITS sequences. Three independent biological replicates (n = 48 samples). Letters indicate statistical significance corresponding to Tukey’s HSD (α = 0.05). NC: normal light condition, LP: low photosynthetically active radiation. **(B)** Variation in B, F, and O community composition explained by “light” (i.e., NC *vs.* LP) in root and matrix samples five weeks post BFO inoculation in the FlowPot system. Constrained analysis of principal coordinates (CAP) was used to quantify the percentage of variance explained by “light” for each condition. *** *P* < 0.001, * *P* < 0.05, ns: not significant. **(C)** Between-group variation in microbial community composition assessed using partial least square discriminant analysis (PLS-DA). Four groups were used as “grouping factor” in the model: NC-Root, NC-matrix, LP-root, and LP-matrix, and associated *P*-value are shown for B (up), F (middle) and O (bottom) community composition. The percentage of variance explained by the grouping factor is indicated along the X and Y axes (B, n = 47 samples, F, n = 48. O, n = 26). **(D)** Normalized relative abundance (RA) of prevalent B strain variants detected in roots across NC and LP conditions. Sample-wise normalized RA was calculated for each strain variant and is depicted in the heatmap next to the taxonomic tree. Only strain variants with an average RA ≥ 0.1% across samples were considered (n = 24 samples). The taxonomic tree was constructed based on a maximum likelihood method using full length 16S rRNA sequences. **(E)** Sample-wise log fold-change in RA measured for each prevalent strain variant between LP and NC in root and matrix samples. Differential RA with statistically significant *P*-values are shown (edgeR, GLM, *P* < 0.05). Strain variants were ordered as depicted in panel D.

### Host prioritizes microbiota-induced growth over defence under LP

We hypothesized that plant responses to BFO commensals and light are interconnected, thereby orchestrating resource investment into shoot under LP. We profiled the root and shoot transcriptomes of BFO-colonized and germ-free *A. thaliana* exposed to LP and NC in the gnotobiotic FlowPot system five weeks post BFO inoculation (**Table S3**). PERMANOVA (**Table S3**) and pairwise correlations among samples (**Figure S3A**) indicated that the presence/absence of the BFO SynCom explained transcriptome differentiation in root samples more than the light condition (microbes: R^2^ = 0.361, *P* < 0.001, light: R^2^ = 0.262, *P* < 0.001; **Table S3**), whereas differentiation in the shoot transcriptome showed the opposite pattern (microbes: R^2^ = 0.224, *P* = 0.003, light: R^2^ = 0.293, *P* < 0.001; **Table S3**). Consistent with a microbiota-root-shoot axis, aboveground responses to light were influenced by BFO commensals (shoot: “microbes:light”, R^2^ = 0.091, *P* = 0.041), whereas belowground responses to microbes were modulated by the light condition (root: “microbes:light”, R^2^ = 0.118, *P* = 0.038) (**Table S3**). Hierarchical clustering of expression profiles of all genes identified as differentially regulated across conditions (|log2FC| ≥ 1, Empirical Bayes Statistics, FDR < 0.05, **Figure S3B**) identified 9 gene expression clusters in root (R1-R9, n = 3,013 genes, **Figure 3A**) and 8 in shoot samples (S1-S8, n = 2,790 genes, **Figure 3B**). Presence of the BFO SynCom triggered light-independent up-regulation of genes involved in ion homeostasis (root, R2), cell differentiation (root, R2), as well as down-regulation of genes involved in response to salicylic acid (SA; root, R9) and anthocyanin biosynthesis (shoot, S6) (**Figures 3A, 3B, S3C** and **S3D, Table S3**). In contrast, LP triggered BFO-independent up-regulation of genes involved in response to jasmonic acid and gibberellin (JA, GA; shoot, S5) as well as downregulation of genes involved in flavonoid metabolic process (shoot, S3) and response to high light intensity (root, R4) (**Figures 3A, 3B, S3C** and **S3D, Table S3**). Furthermore, we identified clusters for which gene expression was modulated by both light and BFO conditions (i.e., R1, R3, R7, R8, S1, S4), likely explaining BFO-mediated host rescue in LP (**Figures 3A** and **3B**). Particularly, genes belonging to clusters R1, R3, and S1 were up-regulated in the presence of the BFO SynCom under NC, but not under LP (**Figures 3A** and **3B**). A substantial fraction of the genes (i.e., 227) did overlap between root clusters R1R3 and the shoot cluster S1 (**Figure 3C**), and GO term analysis revealed consistent enrichments of immune-related processes, including response to chitin, response to SA, JA, and ethylene, response to organonitrogen compounds, or regulation of immune responses, among others (Hypergeometric test with Bonferroni correction, *P* < 0.05, **Figures 3D, S3C** and **S3D, Table S3**). Transcripts with conserved expression patterns in root and shoot primarily encode transcription factors (i.e., WRKY33, WRKY40, MYB15, MYB51, ANAC042, ANAC055), receptor-like protein/kinase (i.e., WAKL10, RLK3, CRK9), ethylene-responsive elements (i.e., ERF6, ERF11), or calmodulin binding proteins (i.e., CBP60G, CML38) (**Figure 3E**). Immune response activation in response to the BFO SynCom under NC was more extensive aboveground than belowground and involved several well-known immune-related genes that act through multiple pathways (**Figure 3E**). These genes encode transcription factors/proteins/enzymes involved in indole glucosinolate biosynthesis (i.e., MYB51, CYP81F2), camalexin production (CYP71B15), SA response (PR1, BGL2, FRK1, EDS5 and PAD4), systemic acquired resistance (SAR; i.e., SARD1, ALD1, AZI1 and DOX1), and to a lesser extend ethylene/JA responses (i.e., PDF1.2b) (**Figure 3E**). Transcriptional down-regulation of these BFO-induced immune responses under LP (S1, R1, R3) were accompanied by transcriptional up-regulation of genes involved in gluconeogenesis in roots, and lipid transport in leaves (R7, S4, **Figure S3E, Table S3**). Our results demonstrate that *A. thaliana* responses to multi-kingdom microbial commensals and light are interconnected along the root-shoot axis, thereby allowing prioritization of microbiota-induced growth over defence responses under LP.

**Figure 3.**
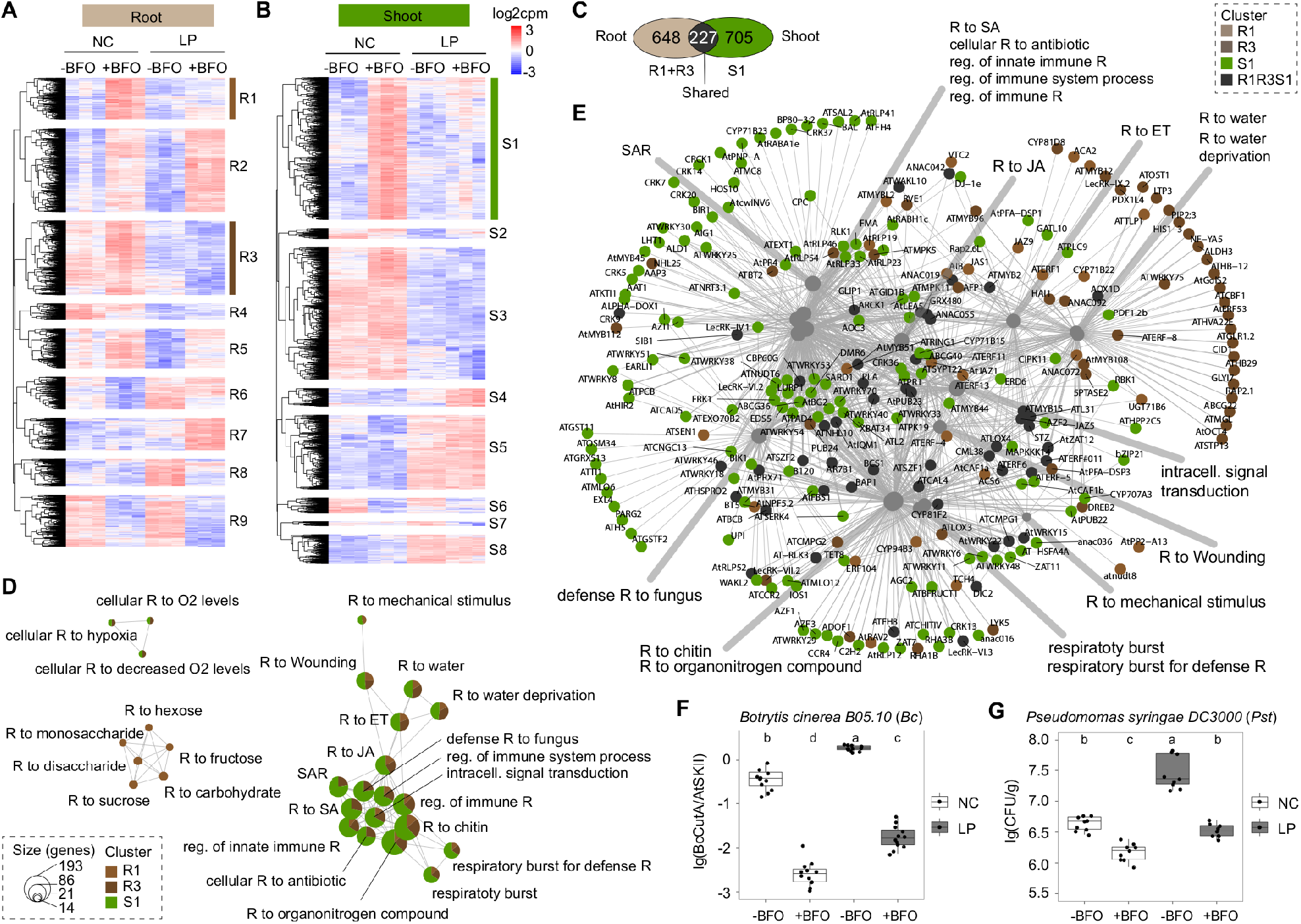
Light- and BFO-mediated transcriptional reprogramming in root and shoot. **(A)** Transcript profiling of 3,013 *A. thaliana* genes significantly regulated in root samples (|log2FC| ≥ 1, Empirical Bayes Statistics, FDR < 0.05) based on all possible pairwise sample comparisons: NC-BFO *vs*. NC+BFO, LP-BFO *vs.* LP+BFO, NC-BFO *vs.* LP-BFO, NC+BFO *vs*. LP+BFO. Over-represented (white to red) and under-represented transcripts (white to blue) are shown across conditions as log2 counts per million (voom-transformed data with limma package in R). The gene set was split into 9 gene expression clusters (R1 to R9). NC: normal light condition, LP: low PAR, -BFO: no microbes, +BFO: with microbes. Three independent biological replicates (n = 12 samples). **(B)** Transcript profiling of 2,790 *A. thaliana* genes significantly regulated in shoot samples (|log2FC| ≥ 1, Empirical Bayes Statistics, FDR < 0.05) based on all possible pairwise sample comparisons (see panel A). The gene set was split into 8 gene expression clusters (S1 to S8). Three independent biological replicates (n = 12 samples). **(C)** Number of genes shared between root clusters R1R3 and shoot cluster S1 or specific to each of the two groups. **(D)** Gene Ontology (GO) term enrichment network depicting the top 12 most significantly enriched GO terms (Hypergeometric test with Bonferroni correction, *P* < 0.05) detected in clusters R1, R3, and S1. Each GO term is represented as a circle and the contribution of each cluster to the overall GO term enrichment is shown. The size of the GO term reflects the number of genes enriched in the GO term. R: response, reg. Regulation, ET: ethylene, SA: salicylic acid, JA: jasmonic acid, SAR: systemic acquired resistance. **(E)** Gene-concept network depicting linkages between genes and associated top 12 most significantly enriched GO terms detected in clusters R1, R3 and S1. Each node represents a gene and is colour-coded according the different cluster names. **(F)** qPCR-based quantification of *Botrytis cinerea B05.10* (*Bc*) growth in *A. thaliana* leaves five days post pathogen inoculation in the FlowPot system. For inoculation, 2 μl droplets containing 1×10^3^ spores were applied to leaves of four-week-old *A. thaliana* grown in the presence/absence of the BFO SynCom under either NC or LP. The relative growth of *Bc* and *A. thaliana* was determined by amplification of the *BcCutinaseA* gene and the *AtSkII*, respectively. Four independent biological replicates (n = 48 plants). Letters indicate statistical significance (ANOVA followed by *post hoc* Tukey’s HSD, α = 0.05). **(G)** Colony count-based quantification of *Pseudomonas syringae pv. tomato* DC3000 (*Pst*) growth in *A. thaliana* leaves five days post pathogen inoculation in the FlowPot system. For inoculation, *Pst* suspension was adjusted to OD = 0.2 and spray-inoculated on leaves of four-week-old *A. thaliana* grown in the presence/absence of the BFO SynCom under either NC or LP. For colony counting, 20 μl dilution series (10^-1^, 10^-2^, 10^-3^, 10^-4^, 10^-5^ with 10 mM Mgcl2) were spotted on NYGA plates, and colonies were counted two days after incubation at 28 °C. Three independent biological replicates (n = 36 plants). Letters indicate statistical significance (ANOVA followed by *post hoc* Tukey’s HSD, α = 0.05).

### Light and root microbiota modulate leaf pathogen resistance

We hypothesised that BFO-triggered aboveground defence responses can protect *A. thaliana* against leaf pathogens in a light-dependent manner. We tested this hypothesis in our gnotobiotic plant system by inoculating leaves of four-week-old *A. thaliana* (NC-BFO, NC+BFO, LP-BFO, LP+BFO) with the necrotrophic fungal pathogen *Botrytis cinerea* B05.10 (*Bc*, droplet inoculation, 1×10^3^ spores) or the biotrophic bacterial pathogen *Pseudomonas syringae pv. tomato* DC3000 (*Pst*, spray inoculation, OD = 0.2). Evaluation of pathogen growth *in planta* by quantitative PCR (*Bc*, **Figure 3F**) or colony counting (*Pst*, **Figure 3G**) revealed major influence of both light and SynCom conditions on disease resistance five days post pathogen inoculation (ANOVA followed by *post hoc* Tukey’s HSD, *P* < 0.05, **Figures 3F** and **3G**). Plants colonized by the BFO SynCom under NC were the most resistant to both leaf pathogens, which is consistent with extensive BFO-triggered immune responses observed in the RNA-Seq data and the presence of putatively protective commensals that ectopically colonize aboveground shoot organs (**Vannier et al., 2019, Figures 3B, 3F** and **3G**). In contrast, germ-free plants facing LP were the most susceptible to both pathogens and failed to mount effective immune responses, indicating that BFO commensals were needed for effective leaf immunity. Although BFO-induced leaf protection against these two pathogens was weakened under LP compared to NC (ANOVA followed by *post hoc* Tukey’s HSD, *P* < 0.05), these plants were still able to limit pathogen proliferation, while massively investing into shoot growth (**Figures 1A**, **3F** and **3G**). Therefore, BFO commensals simultaneously confer aboveground host tolerance to unrelated biotic (microbial pathogens) and abiotic (light limitation) stresses.

### Microbiota-mediated plant rescue depends on several host pathways

BFO-mediated shoot growth promotion under LP likely resulted from complex microbiota-root-shoot signalling mechanisms involving light perception, plant development and immune responses. To identify host components required for prioritization of microbiota-induced growth over defence under LP, we monitored plant growth as well as *Bc* and *Pst* colonization across several mutant plants in our gnotobiotic system (**Figure S4**). These include mutants impaired in SA and JA biosynthesis (SA: *sid2-2*, JA: *dde2-2*; **Wildermuth et al., 2001; von Malek et al., 2002**), JA and GA signalling (JA: *myc2-3/jin1-8* – *jazQ*; GA: *dellaP*; **Shin et al., 2012; Campos et al., 2016; Park et al., 2013**), brassinosteroid signal transduction (*bri1-301*, **Xu et al., 2008**), light perception (*cry1cry2*, **Ahmad et al., 1998**), and indole-3-pyruvic-acid-dependent auxin biosynthesis (*sav3-3*; **Tao et al., 2008**) (**Table S4**). Consistent with previous work (**Xu et al., 2008; Wang et al., 2015; Campos et al., 2016; Pullen et al., 2019**), we observed mutant *versus* Col-0 wild-type plant variation in vegetative growth under control conditions (NC-BFO), with enhanced growth for *dellaP* and *cry1cry2* mutants and reduced growth for *jazQ* and *bri1-301* mutants (**Figures S4A** to **S4J**, Mann-Whitney-U test, *P* < 0.05, statistical analysis not depicted). Irrespective of these differences in growth rate, LP-mediated reduction in canopy size was quantitatively similar across all mutants tested in the absence of the BFO SynCom (0.48 – 0.71-fold decrease in canopy size), except for the *sav3-3* mutant that retained high growth under LP (Kruskal-Wallis with Dunn’s *post hoc* test, **Figure 4A**). The similarity in Col-0 canopy size observed between LP and NC in the presence of BFO (+BFO, ratio LP *vs*. NC close to 1, **Figure 4A**) was largely retained in the *sid2-2, dellaP*, and *sav3-3* mutants, indicating that the corresponding genes are dispensable for BFO-mediated canopy size enlargement under LP (**Figures 4A, S4A** and **S4B**). In contrast, canopy size was dramatically reduced under LP compared to NC for *dde2-2, myc2-3, jazQ, bri1-301* and *cry1cry2* mutants (+BFO, ratio LP vs NC close to 0.5, Mann-Whitney-U test, *P* < 0.01, **Figure 4A, S4A** and **S4B**), indicating that JA biosynthesis/signalling, BR signal transduction, and cryptochromes are needed for BFO-induced growth under LP. Integration of belowground response to microbiota and aboveground response to light therefore involved multiple points of control along the root-shoot axis. In our gnotobiotic system (NC+BFO, NC-BFO, LP-BFO, LP+BFO), we then inoculated leaves of four-week-old *A. thaliana* Col-0 and mutant plants with *Bc* or *Pst*. Consistent with previous work, we observed genotype-specific differences in *Bc* and *Pst* susceptibility phenotypes under control conditions (NC-BFO), with the *dde2-2* and *sid2-2* mutants being the most susceptible and resistant to *Bc*, respectively, and the *sid2-2* and *myc2-3* mutants being the most susceptible and resistant to *Pst*, respectively (**Figures S4K** to **S4N**, **Laluk et al., 2011; Shin et al., 2012; Liu et al., 2017**). For all mutants tested, besides *jazQ (Bc* condition), a significant and consistent increase in *Bc* and *Pst* load was observed in leaves of plants grown in LP compared to NC (*Bc*: 3.70 – 25.69-fold increase in susceptibility, **Figure 4B**; *Pst*: 4.66 – 11.94-fold increase in susceptibility, **Figure 4C**), indicating that LP-mediated reduction in canopy size in the absence of root commensals was also associated with impaired resistance to unrelated leaf pathogens (**Figures 4B** and **4C**). Notably, most of the mutants impaired in BFO-mediated growth rescue in LP showed enhanced resistance towards both *Bc and Pst (myc2-3, jazQ*, and *bri1-301)* or *Pst* only (*cry1cry2*) under LP compared to NC (Mann-Whitney-U test, *P* < 0.05), which is consistent with a prioritization of microbiota-induced defence over growth responses in these mutants. Further analysis of BFO effect size on plant phenotypic traits and pathogen growth under NC and LP (Cohen’s d effect size) supported the role of MYC2 as a potential regulatory node balancing microbiota-induced defence (broad-spectrum) or growth in a light-dependent manner (**Figures 4D** and **4E**).

**Figure 4.**
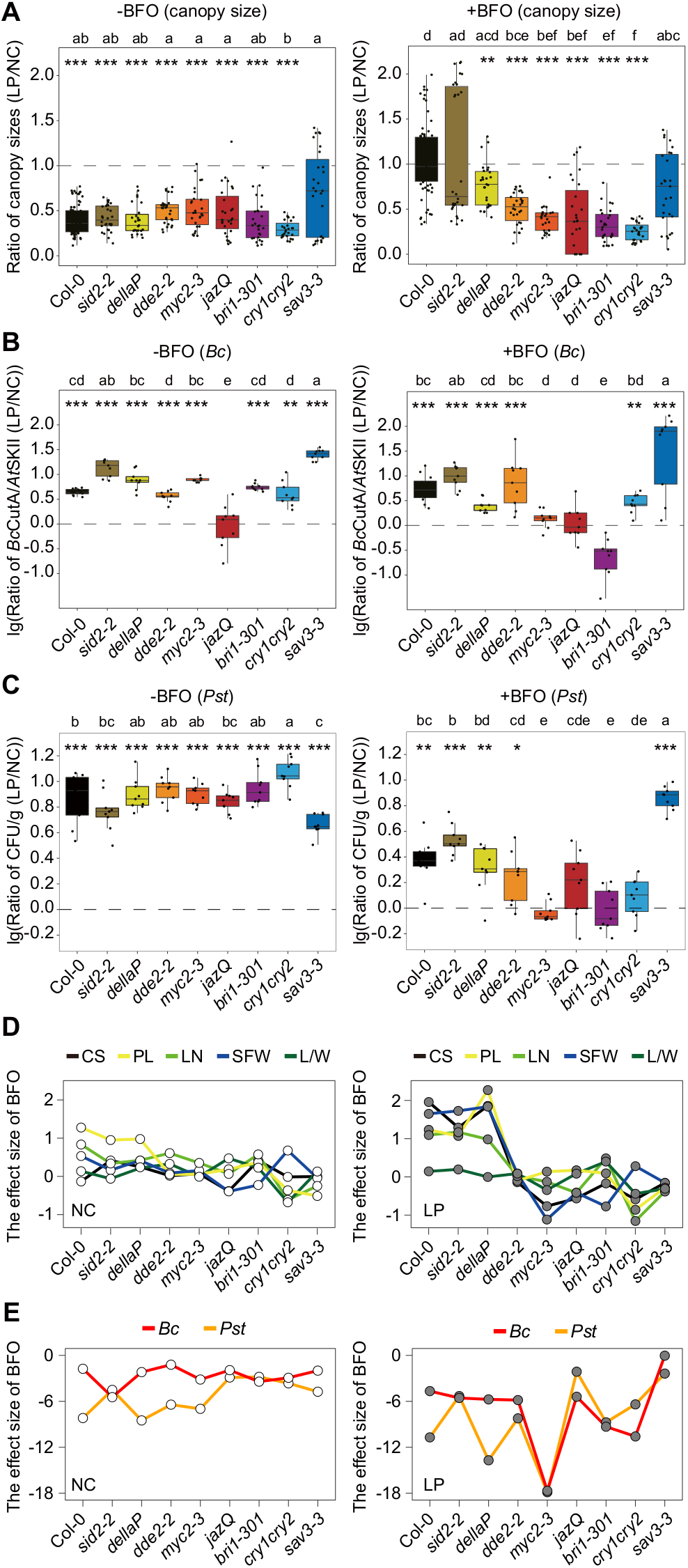
BFO-induced plant growth promotion in LP requires multiple host pathways. **(A)** Differential canopy size between LP and NC for *A. thaliana* Col-0 and eight mutants grown in the FlowPot system under germ-free conditions (-BFO, left) or in the presence of the BFO SynCom (+BFO, right). Ratio in canopy size (five-week-old plants) was computed between LP and NC across three independent biological replicates (n = 1,188, -BFO and +BFO samples). Datapoints below the dashed line showed canopy size decrease in LP compared to NC. Statistical significance across genotypes is indicated with letters (Kruskal-Wallis with Dunn’s *post hoc* test, α = 0.05). Statistical significance between LP and NC is indicated with asterisks (Mann-Whitney U test, *** *P* < 0.001, ** *P* < 0.01, * *P* < 0.05). **(B)** Differential *Botrytis cinerea* B05.10 (*Bc*) growth between LP and NC in leaves of *A. thaliana* Col-0 and eight mutants grown in the FlowPot system (-BFO: left, +BFO: right). *Bc* growth was quantified by qPCR in *A. thaliana* leaves five days post pathogen inoculation. For inoculation, 2 μl droplets containing 1×10^3^ spores were applied to leaves of four-week-old *A. thaliana*. The relative growth of *Bc* and *A. thaliana* was determined by amplification of the *BcCutinaseA* gene and the *AtSkII* genes, respectively. Ratio in *Bc* load was computed between LP and NC across three independent biological replicates for each genotype (n = 336, - BFO and +BFO samples). Datapoints above the dashed showed *Bc* growth increase in LP compared to NC. Statistical significance across genotypes is indicated with letters (ANOVA followed by *post hoc* Tukey’s HSD, α = 0.05). Statistical significance between LP and NC is indicated with asterisks (Mann-Whitney U test, *** *P* < 0.001, ** *P* < 0.01, * *P* < 0.05). **(C)** Differential *Pseudomonas syringae pv. tomato* DC3000 (*Pst*) growth between LP and NC in leaves of *A. thaliana* Col-0 and eight mutants grown in the FlowPot system (-BFO: left, +BFO: right). *Pst* growth was quantified by colony counting in *A. thaliana* leaves five days post pathogen inoculation. *Pst* inoculation was carried out by spaying *Pst* at 0.2 OD in 10 mM MgCl_2_ on leaves of four-week-old *A. thaliana*. Ratio in *Pst* load was computed between LP and NC across three independent biological replicates for each genotype (n = 324, -BFO and +BFO samples). Datapoints above the dashed line showed *Pst* growth increase in LP compared to NC. Statistical significance across genotypes is indicated with letters (ANOVA followed by *post hoc* Tukey’s HSD, α = 0.05). Statistical significance between LP and NC is indicated with asterisks (Mann-Whitney U test, *** *P* < 0.001, ** *P* < 0.01, * *P* < 0.05). **(D)** Effect size of BFO on shoot morphological traits of *A. thaliana* Col-0 and eight mutants grown in the FlowPot system under NC (left) and LP (right). Effect size was computed between -BFO and +BFO conditions for each light condition by adjusting the calculation of the standard deviation with weights of the sample sizes (Cohen’s d). Different line colors reflect different shoot morphological traits. CS: canopy size, PL: petiole length (mean/plant), LN: leaf number, SFW: shoot fresh weight, L/W: leaf length/width ratio. **(E)** Effect size of BFO on *Botrytis cinerea* B05.10 (*Bc*) growth (red line) and *Pseudomonas syringae pv. tomato* DC3000 (*Pst*) growth (orange line) in leaves of *A. thaliana* Col-0 and eight mutants grown in the FlowPot system under NC (left) and LP (right). Effect size was which the different mutants were grown in the gnotobiotic system under NC and LP to simultaneously monitor BFO assemblages in roots and surrounding peat matrix five weeks post BFO inoculation. PERMANOVA confirmed the effect of the light condition on B community composition in the roots, but not in matrix samples (root, “light”: R^2^ = 0.014, *P* = 0.024) and revealed that B community differentiation was more extensively shaped by “genotype” than by “light” (root, “genotype”: R^2^ = 0.117, *P* < 0.001), which was validated by CAP (**Table S5** and **Figure S5A**). CAP analysis constrained by “light” for each genotype indicated that light-mediated differentiation in B community composition was greater for *jazQ* (12.8%, *P* = 0.004), *dde2-2* (8.07%, *P* = 0.0265), *myc2-3* (7.87%, *P* = 0.005), *bri1-301* (6.26%, *P* = 0.049), than for WT (3.35%, *P* = 0.044), and not significant for the other mutants (**Figure S5B**). Pairwise-enrichment tests conducted between LP and NC conditions for each genotype (edgeR, generalized linear model, *P* < 0.05, **Figure 5A**) validated the increase in RA of root-associated *Pseudomonas* observed in WT, and revealed mutant-specific differences in abundance profiles at the strain (**Figure 5A**) and class (**Figure S5C**) levels. The lack of enrichment of *Pseudomonas* in roots of most of the mutants that lack BFO-mediated growth rescue in LP (**Figure 5A**) prompted us to first test the extent to which the seven *Pseudomonas* isolates used in our SynCom were needed for BFO-mediated growth rescue in LP. Remarkably, a B community lacking these seven strains (i.e. 177 members) was no longer able to rescue plant growth under LP, whereas a mini SynCom consisting of the seven *Pseudomonas* strains alone showed an intermediate, yet not significant

### A link between differential canopy size and root microbiota composition under LP

We asked whether differentiation in B community composition in the roots of WT plants was altered in the different mutants and could correlate with aboveground change in canopy size. We took advantage of the previous experiment in growth rescue (**Figure 5B**, Col-0 background). These results demonstrate that *Pseudomonas* root commensals are necessary, although not sufficient, for BFO-mediated growth rescue under LP. To further test for potential associations between aboveground canopy phenotypes and belowground B community composition, we calculated canopy size variation for each mutant between LP-BFO and LP+BFO conditions, and asked whether these quantitative differences were significantly linked to corresponding root microbiota composition under LP. Using a linear regression model, we observed a significant link between BFO-induced canopy size across mutants and B community composition in roots (coordinate on PCOA axis 1, 46.9% of the variance in Bray-Curtis dissimilarities), which explained 47.7% of the variation in B community composition along PCoA1 (R^2^ = 0.4776, F_17_ = 8.314, *P* = 0.02353, **Figure 5C**). We then divided the genotypes into two groups based on the BFO-mediated growth induction (rescued: BFO-induced growth under LP; not rescued: lack of BFO-induced growth under LP) and validated that belowground change in B community composition can be used to discriminate those groups, with a mean classification error of 20.4% (PLS-DA, *P* < 0.001, **Figure 5D**). To identify the strain variants that allow the best discrimination of the two groups based on variations of their RA in LP, we trained a Support Vector Machines classifier (SVM) with Recursive Feature Elimination, and identified a set of 37 strain variants that were sufficient to accurately predict the phenotype group: R^2^ = 0.83 (**Figure S5D, Table S5**). Using RA data of these 37 strain variants as an input for PLS-DA, we greatly improved the model quality (PLS-DA, *P* < 0.001, classification error = 8.1%, **Figure 5E**), whereas a similar analysis with the strain variants not selected by the SVM classifier (n = 68) could no longer discriminate the two groups (PLS-DA, *P* > 0.05, classification error > 40%, data not shown). Taken together, our results suggest that plant growth phenotypes under LP and belowground B community composition are linked and that *Pseudomonas* root commensals are key community members necessary for BFO-mediated growth rescue.

**Figure 5.**
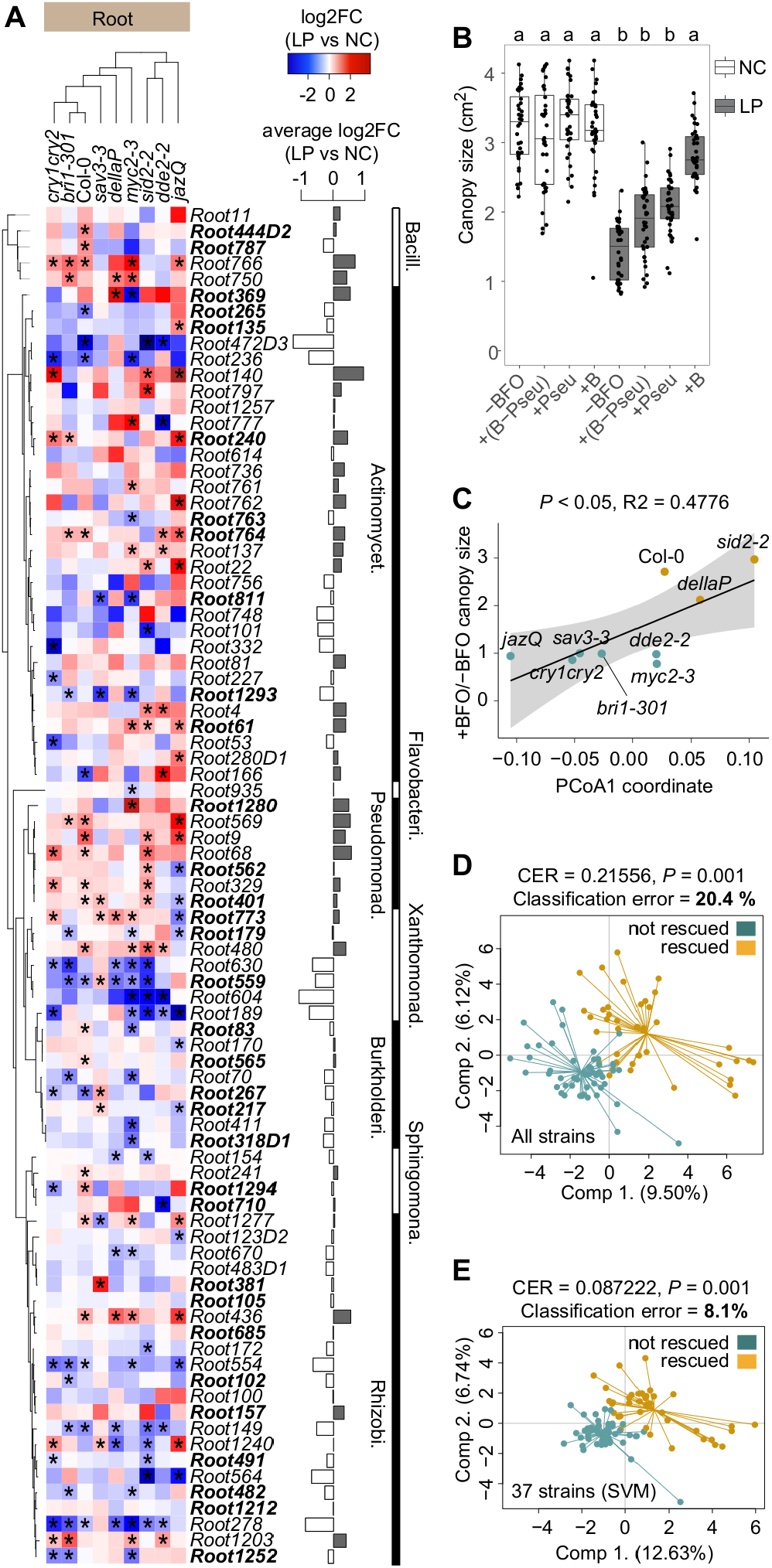
Link between genotype-dependent B community shifts and BFO-induced growth in LP. **(A)** Phylogeny-based heatmap showing differential relative abundance (Log2FC, root samples) between LP and NC for each strain variant across different *A. thaliana* genotypes. The Maximum likelihood tree includes only strain variants (n = 85) that were consistently present across genotype samples (number of samples per genotype: n = 9, except *jazQ*: n = 6 and Col-0: n = 21). Differences between LP and NC were calculated based on mean values and are shown as a gradient from blue (LP < NC) to red (LP > NC). Taxonomy is shown at the class level with alternating white and black colours. Samples (i.e., genotypes) were ordered based on hierarchical clustering. Significant differences in RA between LP and NC are indicated by asterisks (edgeR, GLM, *P* < 0.05). The barplot next to the heatmap represents the mean difference in RA between LP and NC measured across all genotypes. Strain variants highlighted in bold correspond to those identified through the Support Vector Machines classifier. **(B)** Canopy size (in cm^2^) of five-week-old *A. thaliana* grown in the FlowPot system in the absence (-) or presence (+) of a synthetic microbial community composed of either B without *Pseudomonas* strains (+(B-Pseu)), or *Pseudomonas* strains only (+Pseu), or B (+B) under NC (white) and LP (grey). Three independent biological replicates (n = 288 plants). Letters indicate statistical significance corresponding to Kruskal-Wallis with Dunn’s *post hoc* test (α = 0.05). **(C)** Linear regression between BFO-induced canopy size in LP and bacterial community composition. BFO-induced canopy size was calculated as a ratio between the canopy size under LP in +BFO and mean canopy size of the respective mutant under LP in - BFO. Coordinates on the first axis of a PCOA based on bray-curtis dissimilarities between samples was used as a proxy of bacterial community composition. P-value and R^2^ obtained with ANOVA are indicated in the figure. **(D)** Partial least square discriminant analysis (PLS-DA) testing the differentiation between rescued and non-rescued plant phenotypes in LP based on the composition of the bacterial communities across all mutants (i.e., strains RA; 109 strains, samples n = 90). Quality parameters of the analysis (P-value, CER and mean misclassification rate) are indicated in the figure. **(E)** SVM-guided Partial least square discriminant analysis (PLS-DA) testing the differentiation between rescued and non-rescued plant phenotypes in LP based on the composition strains RA using the subset of strains identified in the System Vector Machine (SVM) (37 strains, samples n = 90). Quality parameters of the analysis (P-value, CER and mean misclassification rate) are indicated in the figure.

### Priority to microbiota-induced growth over defence under LP requires MYC2

Given the fact that MYC2 is a central node controlling the crosstalk between JA and other phytohormone signalling pathways (i.e., GA, SA, ABA, IAA), and regulating responses to light and the circadian clock (**Kazan and Manners, 2013**), we hypothesized that this transcription factor might coordinate prioritization of microbiota-induced growth over defence under suboptimal light conditions. Importantly, impaired growth and increased resistance to *Bc* observed in the *myc2-3* mutant were quantitatively similar in the independent *myc234* triple mutant (**Figure S4O**), and reverted in the *jin1-8* pMYC2:MYC2-FLAG line in which a MYC2-FLAG fusion protein is expressed under control of the native MYC2 promoter (MYC2-FLAG, *myc2* background, **Hou et al., 2010**) (**Figures 6A**). We also demonstrated that alleviation of EODFR-mediated decrease in canopy size by BFO also required MYC2, illustrating that MYC2-dependent rescue of plant growth by BFO is robust across multiple light-limiting conditions (**Figure 6A**). Western blot assays revealed that the expression level of MYC2 in root and shoot samples was modulated by BFO under LP but not under NC (MYC2-FLAG line), with expression levels in the shoot matching canopy size phenotypes (**Figures 6B** and **S6A**). We profiled the root and shoot transcriptomes of BFO-colonized and germ free *myc2-3* and MYC2-FLAG lines exposed to LP and NC in the gnotobiotic FlowPot system five weeks post BFO inoculation. Remarkably, presence/absence of the BFO SynCom explained transcriptome differentiation in shoot samples more than light in the *myc2-3* mutant (**Figure S6B**), which was validated by PERMANOVA (*myc2-3*, microbes: R^2^ = 0.394, *P* < 0.001, light: R^2^ = 0.179, *P* = 0.006; MYC2-FLAG, microbes: R^2^ = 0.085, *P* = 0.078, light: R^2^ = 0.439, *P* < 0.001, **Table S6**). These results suggest attenuation of shoot response to light and/or exacerbation of shoot response to microbes in this mutant. Mutation of MYC2 led to dramatic shifts in the root and shoot transcriptome, with 5,231 and 5,038 genes differentially regulated in *myc2-3* compared to MYC2-FLAG in root and shoot, respectively (referred to as *Myc2*-Differentially Expressed or MDEs, |log2FC| ≥ 1, Empirical Bayes Statistics, FDR < 0.05, **Figures 6C, S6C** and **S6D**). A large fraction of these MDEs were previously reported as direct targets of MYC2 (MDEs root: 52%, MDEs shoot: 58%, **Figure 6D** and **S6E**, **Van Moerkercke et al., 2019; Wang et al., 2019; Zander et al., 2020**). Hierarchical clustering of gene expression profiles of MDEs revealed 8 gene expression clusters for both root and shoot samples. In roots, GO terms enrichment analyses of MDEs for each cluster revealed that processes related to ion homeostasis and circadian rhythm (MDER6), response to sugar (MDER2), defence (MDER5), photosynthesis and response to light (MDER6), or cytokinesis processes (MDER6) were modulated by MYC2 across the different conditions (**Figure S6F**). In shoots, terms related to photosynthesis (MDES1), defence (MDES2), cytokinesis (MDES3), secondary metabolite biosynthesis (MDES4), response to JA (MDES5), or sugar biosynthesis (MDES7) were also altered in the *myc2-3* mutant (**Figure S6F**). Notably, clusters MDES2 and MDES7 contained GO terms for which the most significant enrichments were observed and genes in these clusters were either induced (MDES2) or repressed (MDES7) in the *myc2-3* mutant compared to MYC2-FLAG under LP in the presence of BFO (**Figures 6C** and **S6F**). Therefore, MYC2-dependent transcriptional reprogramming between these two clusters likely explained the lack of BFO-mediated growth rescue under LP in the mutant. Inspection of the GO terms and associated genes revealed that BFO-triggered shoot defence responses were retained in the *myc2-3* mutant under LP (MDES2), with 28% of the genes shared between this cluster and the previously identified cluster S1 (**Figures 6E** and **6F**, see also **Figure 3B**).

**Figure 6.**
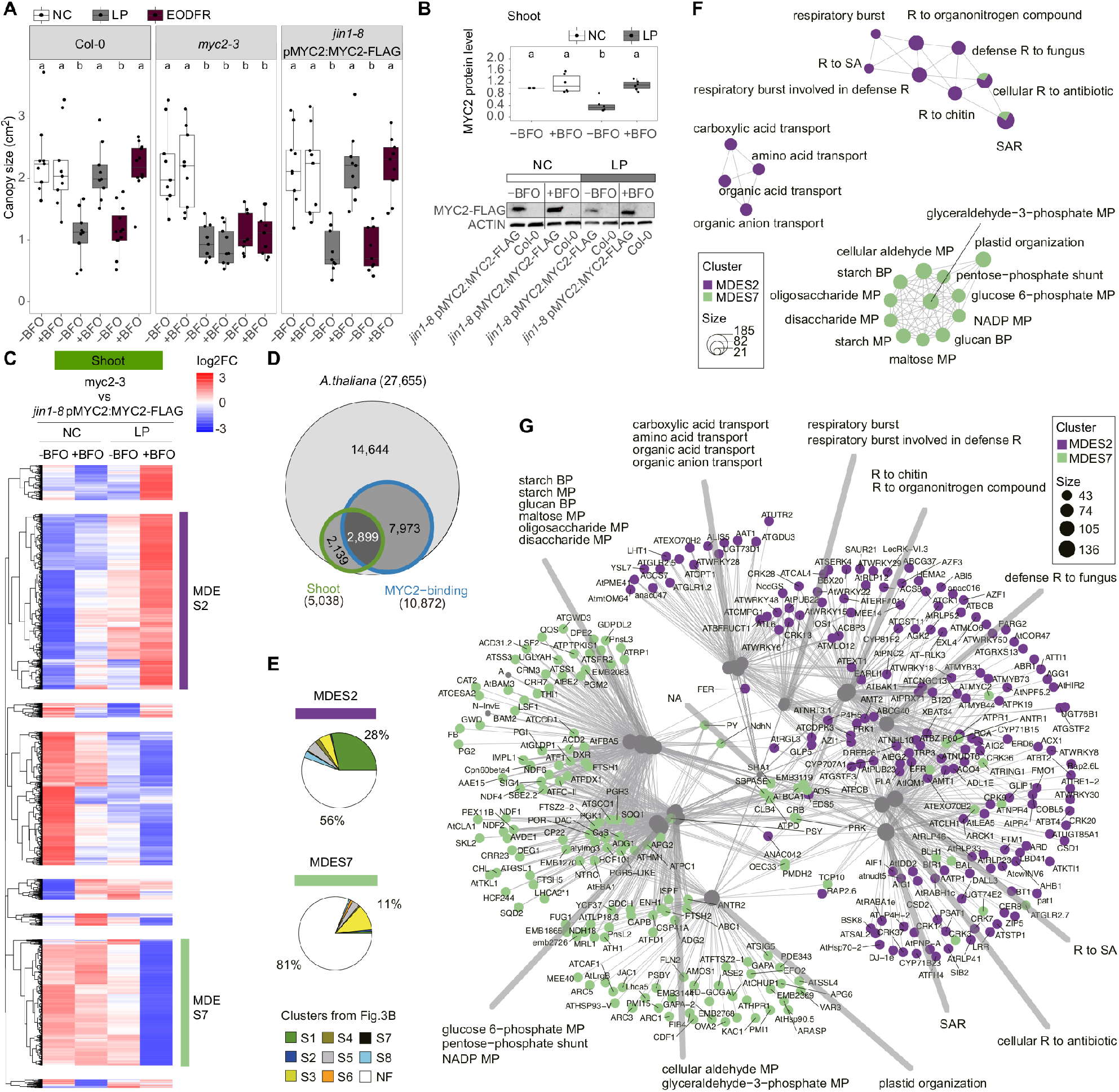
MYC2-dependent prioritization of BFO-induced growth over defence in LP. **(A)** Canopy size (in cm^2^) of five-week-old WT (Col-0), mutant (*myc2-3*), and complemented (*jin1-8* pMYC2:MYC2-FLAG) *A. thaliana* grown in the FlowPot system in the absence (-) or presence (+) of the BFO SynCom under either normal light (NC, white), low photosynthetically active radiation (LP, grey), or end-of-day far-red (EODFR, dark red). Three independent biological replicates (Col-0: n = 60 plants, *myc2-3:* n = 54 plants, *jin1-8 pMYC2:MYC2-FLAG:* n = 54 plants). Letters indicate statistical significance (Kruskal-Wallis with Dunn’s *post hoc* test, α = 0.05). **(B)** Quantified MYC2 protein levels in shoots of five-week-old *jin1-8* pMYC2:MYC2-FLAG line grown in the FlowPot system in the absence (-) or presence (+) of the BFO SynCom under either NC (white) or LP (grey). Col-0 shoots under corresponding conditions were as control samples. MYC2 protein levels first were quantified by ACTIN then normalized to NC-BFO condition. Three independent biological replicates including six independent replicates of immunoblots (n = 48 samples). Letters indicate statistical significance (Kruskal-Wallis with Dunn’s *post hoc* test, α = 0.05). One immunoblots replicate was shown on the bottom. **(C)** Transcript profiling of 5,038 *A. thaliana* genes significantly regulated in shoot samples between *myc2-3* mutant and *jin1-8* pMYC2:MYC2-FLAG line (|log2FC| ≥ 1, Empirical Bayes Statistics, FDR < 0.05). Over-represented (white to red) and under-represented transcripts (white to blue) are shown across conditions as mean the differential gene expression between *myc2-3* and *jin1-8* pMYC2:MYC2-FLAG (log2 fold-change of counts per million (TMM-normalized voom-transformed data with limma package in R)). The gene set was split into eight major MYC2 differentially-expressed gene expression clusters in shoots, labelled MDES1 to MDES8. NC: normal light condition, LP: low PAR, -BFO: no microbes, +BFO: with microbes. Three independent biological replicates (n = 24 samples). **(D)** Overlap between the number of MDE genes identified in leaves and all MYC2 target genes identified by ChipSeq experiments in three independent studies (**Van Moerkercke et al., 2019; Wang et al., 2019; Zander et al., 2020**). **(E)** Percentage of genes shared between shoot clusters MDES2 (top) and MDES7 (bottom) and shoot clusters (S1 to S8) previously defined in **Figure 3B**. **(F)** Gene Ontology (GO) term enrichment network depicting the top 12 most significantly enriched GO terms (Hypergeometric test with Bonferroni correction, *P* < 0.05) detected in clusters MDES2 and MDES7. Each GO term is represented as a circle and the contribution of each cluster to the overall GO term enrichment is shown. The size of the GO term reflects the number of genes enriched in the GO term. R: response, SA: salicylic acid, SAR: systemic acquired resistance, BP: biosynthetic process, MP: metabolic process. **(G)** Gene-concept network (cnetplot function in R) depicting linkages between genes and associated top 12 most significantly enriched GO terms detected in clusters MDES2 and MDES7. Each node represents a gene and is colour-coded according the different cluster names.

Consistent with the high levels of free SA (i.e., active) measured in leaves of the *myc2*-3 mutant under LP (**Figure S6G**), these responses involved primarily SA-and/or SAR-related genes (i.e., PR1, BGL2, FRK1, EDS5, SARD1, AZI1 and FMO1), indicating that BFO-triggered immunity in leaves remained activated under LP in this mutant (**Figure 6G**). In contrast, genes involved in starch and sugar metabolic processes were down-regulated under the same condition, illustrating that priority to defence in shoot of the *myc2*-3 mutant was associated with altered sugar metabolic processes. These genes encode enzymes primarily involved in starch biosynthesis (i.e., SS1, SS3), starch accumulation (i.e., ADG1, PGI), or starch breakdown (i.e., LSF1, LSF2, PTPKIS1, BAM3) in shoots (**Figure 6G**). Our results indicate that the trade-off between starch/carbon metabolism and defence in shoots is modulated by microbiota and light in a MYC2-dependent manner.

## DISCUSSION

Reminiscent to the bidirectional communication mechanisms described along the microbiota-gut-brain axis in animals (**Carabotti et al., 2015; Dinan and Cryan 2017, Stassen et al., 2020**), we report here that aboveground stress responses in plants can be orchestrated through long-distance communication with belowground commensals. We demonstrate that change in aboveground light can cascade along the shoot-root axis to alter the composition of root commensal communities. Reciprocally, the presence of BFO commensals triggered leaf activation of defence or growth responses in a light-dependent manner. Integration of these responses to microbes and light along the root-shoot axis dictates the trade-off between growth and defence, thereby boosting plant growth under suboptimal light conditions (**Figure 7**).

**Figure 7.**
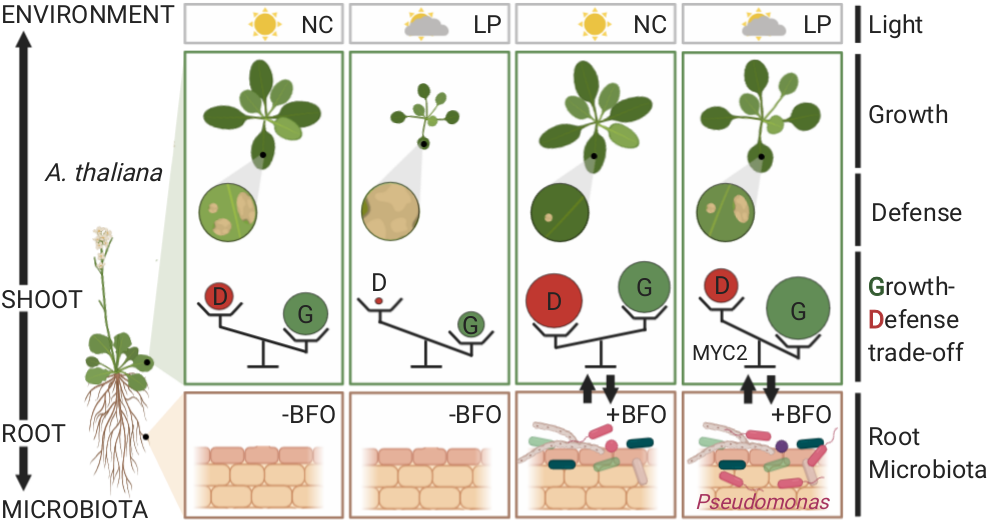
Model for root microbiota-induced growth-defence trade-off between NC and LP. Graphical summary illustrating the bidirectional communication mechanisms along the microbiota-root-shoot-environment axis in *A. thaliana*. In the absence of BFO commensals, plants prioritize growth over defence, especially under LP. However, suboptimal light conditions restrict both growth and defence responses, leading to small plants that are highly susceptible to leaf pathogens. In the presence of BFO root commensals under NC, extensive activation of immunity response was observed in leaves, thereby effectively protecting leaves against microbial pathogens. The BFO SynCom also promotes growth responses under this condition, which likely compensates the fitness cost associated with this elevated immune status. Under suboptimal light conditions, root microbiota-induced systemic immune responses in leaves are shut down in a MYC2-dependent manner, thereby giving priority to microbiota-induced growth. However, although these plants massively invest into shoot growth in LP, they remain more resistant to leaf pathogens than corresponding germ-free control plants.

Over the course of evolution, roots of land plants have continuously interacted with multi-kingdom microbial commensals (**Hassani et al., 2018; Getzke et al., 2019; Martin et al., 2017**). Consequently, microbial molecules in the rhizosphere (i.e., hormones, microbe-associated molecular patterns, volatile organic compounds, lipo-chitooligomers) have been shown to serve as developmental signals for plant growth and immune system maturation (**Ortiz-Castro et al., 2009; Selosse et al., 2014; Vannier et al., 2019**). However, whether microbial signals belowground can modulate aboveground stress responses and *vice versa* remains poorly understood. Bidirectional communication mechanisms between shoot and root organs have been previously described in the context of aboveground biotic stresses. Leaf colonization by microbial pathogens or herbivores resulted in shifts in the rhizosphere microbiota through host-induced modulation of root exudation profiles (**Rudrappa et al., 2008; Yuan et al., 2018; Hu et al., 2018**). Using manipulation experiments, these pathogen/herbivore-induced shifts in rhizosphere commensal communities were shown to be the direct cause protecting the next plant generation through the promotion of systemic defence responses (**Berendsen et al., 2018; Yuan et al., 2018; Hu et al., 2018**). Therefore, modulation of the rhizosphere microbiota via leaf pathogen-induced change in root exudation profiles appears to dictate survival and performance of the offspring (i.e. cry-for-help hypothesis). Although we identified a clear link between differential canopy size across mutants and bacterial community composition under LP, it remains difficult to experimentally test whether these shifts have any physiological relevance for BFO-induced growth-defence trade-off between NC and LP. Using drop-out experiments, we nonetheless observed that the *Pseudomonas* isolates that were enriched in roots of plants facing LP were necessary for the growth rescue phenotype, suggesting that recruitment of these commensals might further enhance prioritization of BFO-induced growth over defence responses under LP.

Induction of systemic defence responses, including pathogen-triggered systemic acquired resistance (SAR, SA-dependent) and commensal-triggered induced systemic resistance (ISR, JA- and ET-dependent), has been extensively reported in response to specific commensal or pathogenic microbes (**Chanda et al. 2011; Cecchini et al. 2015; Zamioudis et al., 2015; Vishwanathan et al. 2020; Wenig et al., 2019**). Here, we showed that root colonization by a multi-kingdom consortium of microbes originally isolated from roots of healthy plants, triggered aboveground induction of defence responses effective against *Bc* and *Pst*. Since ectopic leaf colonization by few root microbiota members was also noted, we cannot exclude the possibility that these defence responses also include local immune responses activated in response to these ectopic colonizers. The observed defence responses induced by BFO commensals in leaves of WT (cluster S1, NC+BFO) and *myc2-3* (cluster MDES2, LP+BFO) plants resemble more stereotypical SAR than ISR responses (**van Wees et al., 2000**), which is consistent with the fact that ISR responses were previously found to be abolished in the *myc2-3* mutant (**Pozo et al., 2008**). Induction of immune responses is costly for plants (**Walters and Heil 2007, Bernsdorff et al., 2016**) and, consistent with our data (**Figure 6**), these responses are known to be associated with the down-regulation of genes involved in chloroplast functions, including photosynthetic light reactions, the Calvin cycle, photorespiration, and starch metabolism (**Gruner et al., 2013, Bernsdorff et al., 2016, Schwachtje et al., 2018**). As chloroplasts are central integrators of multiple functions linked to photosynthesis, defence, and development (i.e., **Chanda et al., 2011, Cecchini et al., 2015, Serrano et al., 2016**), complex metabolic trade-offs in these organelles likely dictate plant investment into microbiota-induced growth or microbiota-induced defence responses.

A direct link between induced defences and light conditions is supported by previous work showing an absolute requirement of phytochromes, but not cryptochromes, for biological induction of SAR (**Zeier et al., 2004; Griebel and Zeier 2008**). Since phytochrome-mediated light signalling is interconnected with JA signalling, likely via MYC2 (**Robson et al., 2010; Gangappa et al., 2010; Prasad et al., 2012; Campos et al., 2016**), it has been suggested that MYC2 might orchestrate the regulation of plant growth and development by light quality (**Kazan and Manners 2013**). Our results are consistent with this hypothesis and suggest that Lp-dependent down-regulation of microbiota-induced defence in leaves is orchestrated through MYC2 via a cross talk between photoreceptor signalling and defence signalling. Our observation that MYC2 is needed for LP-triggered reduction of immune activity against *Pst* and *Bc* in the presence of BFO commensals but not under axenic conditions indicates that this MYC2-dependent growth-defence trade-off requires signals from microbial commensals. This result further supports the hypothesis that, over evolutionary time, direct integration of microbial and environmental cues by plants has been key for plant adaptation to environmental constraints.

Our results suggest that interference of MYC2 (JA signalling pathway) with light, SA/SAR and GA signalling pathways is key to prioritize investment in shoot growth over shoot defence under LP. In our gene expression network (**Figure 6**), we identified few genes connecting the clusters MDE_S2 (growth cluster) and MDE_S7 (defence cluster). Remarkably, these MDEGs include EDS5 (**Serrano et al., 2013**), a transporter required for SA export from chloroplast and needed for SA signalling (induced in *myc2-3* in LP); AOS (**Sanders et al., 2000**), a key chloroplastic enzyme needed for JA biosynthesis (repressed in *myc2-3* in LP); and RGL3 (**Dill and Sun 2001**), a repressor of GA responses (induced in *myc2-3* in LP). As these genes are well-known MYC2 target genes (**Mine et al., 2017; Wild et al., 2012; Van Moerkercke et al., 2019; Zander et al., 2020**), it is conceivable that differential regulation of the expression of these genes by MYC2 orchestrates a complex cross talk between JA, SA, and GA to either prioritize microbiota-induced growth or defence according to the surrounding light conditions. Importantly, this survival trade-off might not necessarily occur in plant species that naturally grow at high density or under suboptimal light conditions, as exemplified by the concomitant activation of defence and shade avoidance responses observed in *Geranium robertianum* (**Gommers et al., 2017**).

Taken together, our data suggest that plant growth and survival in nature likely depends on the ability of these sessile organisms to utilize belowground microbial signals to either prioritize growth or defence depending on light quantity/quality perceived by leaves. Therefore, phenotypic plasticity and aboveground stress responses in plants can be governed by microbial root commensals.

## ACKNOWLEDGEMENTS

This work was supported by funds to S.Ha from a European Research Council starting grant (MICRORULES 758003), the Max Planck Society, as well as the Cluster of Excellence on Plant Sciences (CEPLAS), and the ‘Priority Programme: Deconstruction and Reconstruction of the Plant Microbiota (SPP DECRyPT 2125)’, both funded by the Deutsche Forschungsgemeinschaft. S.Ho. salary was initially covered by a scholarship provided by the China Scholarship Council (CSC Student ID 201604910525), and later on by the ERC MICRORULES grant. We thank H. Frerigmann for sharing the mutant *dellaP* and *jazQ* seeds, T. Nobori for sharing the mutant *dde2-2* and *sid2-2* seeds, J. Qiu for sharing the *myc2-3* mutant, *myc234* triple mutant and *jin1-8* pMYC2:MYC2-FLAG line seeds, Y. Belkhadir for sharing the mutant *bri1-301* and *sav3-3* seeds, and Z. Hu for sharing the *cry1cry2* mutant seeds. We thank R. Birkenbihl for providing the *Botrytis cinerea* B05.10 strain, and the Parker lab for providing *Pseudomonas syringae* pv. *tomato* DC3000 strain. We thank H. Gao for suggestions of low PAR and EODFR conditions. Finally, we thank Prof. Paul Schulze-Lefert, Dr. Ruben Garrido-Oter, Dr. Ryohei Thomas Nakano, Dr. Kenichi Tsuda, Prof. Maria von Korff Schmising and Rozina Kardakaris for providing helpful comments regarding the manuscript or during departmental seminars and thesis advisory committee meetings.

## AUTHOR CONTRIBUTION

S.Ha. initiated, coordinated and supervised the project. S.Ho performed all the experiments and analysed the data, except community profiling data that were primarily analysed by T.T. and N.V. F.M. provided scripts and expertise linked to support vector machines. B.P. measured shoot traits parameters and maintained microbial culture collections. J.Z. quantified phytohormone levels from plant root and leaf samples. S.Ha. wrote the manuscript, with input from S.Ho, T.T., and N.V.

## MATERIALS AND METHODS

### Microbial strains and plant model

The bacterial, fungal and oomycetal strains used in this study have been previously reported (**Bai et al., 2015, Durán et al., 2018**) and are summarized in **Table S1**. *Pseudomonas syringae* pv. *tomato* DC 3000 and *Botrytis cinerea* B05.10 (a benomyl derivative of the strain SAS56) were used as model pathogens for pathogen assays in *A. thaliana* (**Whalen et al., 1991, Liu et al., 2017**). Bacterial strains were routinely cultured at 25°C in liquid 50% TSB medium (Sigma-Aldrich, USA) and stored at −80°C in 25% glycerol. Fungal strains were cultured at 20°C in solid PGA media (Sigma-Aldrich, USA) and agar plugs containing mycelia were stored at −80°C in 30% glycerol. Oomycetal strains were continuously propagated in solid PGA media. *Pseudomonas syringae* pv. *tomato* DC 3000 was cultured at 28°C in NYGA medium and stored at −80°C in 7% Dimethyl sulfoxide (DMSO). *Botrytis cinerea* B05.10 spores were obtained from −80°C glycerol stocks in which the concentration was adjusted to 10^7^ spores ml^-1^ in Vogelbuffer (in 1L: 15 g sucrose, 3 g Na-citrate, 5 g K2HPO4, 0.2 g MgSO4 7H2O, 0.1 g CaCl2 2H2O, 2 g NH4NO3). *A. thaliana* Col-0 and the mutants used in this study are provided in this section and **Table S4**. *A. thaliana* Col-0 wild-type (N60000) was obtained from the Nottingham Arabidopsis Stock Centre (NASC). The *dellaP* mutant (*della* pentuple) was previously generated by crossing *ga-28, rgl1-SK62, rgl2-SK54, rgl3-3*, and introgressed *gai-t6* (**Park et al., 2013**). The *jazQ* mutant (*jaz* quintuple) was previously obtained from T-DNA insertion mutants of *jaz1-2, jaz3-4, jaz4-1, jaz9-4* and *jaz10-1* (**Campos et al., 2016**). The mutants *dde2-2* (CS65993) (**von Malek et al., 2002**), *sid2-2* (CS16438) (**Wildermuth et al., 2001**), *myc2-3* (salk_061267) (**Shin et al., 2012**), *sav3-3* (**Tao et al., 2008**), *bri1-301* (**Xu et al., 2008**), *myc234* (**Fernandez-Calvo et al., 2011**), *cry1-304 cry2-1* (**Mockler et al., 1999**), *jin1-8* pMYC2:MYC2-FLAG (**Hou et al., 2010**) were previously reported. Seeds of *A. thaliana Col-0* and mutant plants were surface-sterilized in 70% ethanol for 18 min followed by a brief wash with 100% ethanol (1 min). Seeds were dried out under sterile bench conditions, and were incubated for two days at 4°C in the dark. Individual seeds were sown onto the surface of FlowPots (**Kremer et al., 2018**) by pipetting one seed at a time. Before seed sowing, FlowPots were inoculated with half-strength Murashige and Skoog medium (MS, without sucrose, pH 5.5, Sigma-Aldrich, USA) with or without microbial commensals. Combined boxes containing FlowPots with sterile or colonized plants were incubated under short-day conditions at 21°C with three light conditions (NC: PPFD 62.35 μmol/m^2^/s, LP: PPFD 27.91 μmol/m^2^/s, EODFR: 15min far-red light (740 nm) treatment at the end of the day) (10 hr) and at 19°C in the dark (14 hr).

### Microbiota reconstitution experiments in the FlowPot system

Bacterial strains were cultivated in 96-deep-well plates containing 400 μl of 50% TSB (Tryptic Soy Broth, Sigma) with 180 rpm shaking speed for seven days at 25°C and subsequently pooled at equal volume ratios. This bacterial pool was centrifuged at 4,000 xg for 10 min and re-suspended in 10 mM MgCl_2_ to remove residual media and bacteria-derived metabolites. Prior to inoculation, OD600 was adjusted to 0.5. Individual fungal and oomycetal strains were cultivated on solid PGA medium for fourteen days. 100 mg fungal and 40 mg oomycetal mycelium was harvested for each strain and aliquoted into 2 ml Eppendorf tubes containing 1 ml MgCl_2_ and one sterile stainless-steel bead (3.2 mm). The mycelium was subsequently crushed with a paint shaker (SK450, Fast & Fluid Management, Sassenheim, Netherlands) for 10 min. Fragmented fungal or oomycetal mycelia were then pooled at equal volume ratios at a concentration of 100 mg/ml for fungi and 40 mg/mL for oomycetes. For the microbial ratio experiment, either 2 ml (high concentration, H) or 0.2 ml (low concentration, L) of the bacterial suspension (OD600 = 0.5, see above), 1 ml (H) or 0.1 ml (L) of the fungal suspension (100mg/mL, see above), and 1 ml (H) or 0.1 ml (L) of the oomycetal suspension (40 mg/mL, see above) was transferred into a falcon containing 50 ml of MS medium without sucrose, which was used to repopulate sterile peat in the FlowPot. For all the other experiments in the FlowPot system, 0.2 ml (L) of the bacterial suspension (OD600 = 0.5), 0.1 ml (L) of the fungal suspension (100 mg/mL), and 0.1 ml (L) of the oomycetal suspension (40mg/mL) were used. Procedures for setting up the gnotobiotic FlowPot system were carried out as previously described (**Kremer et al., 2018, Durán et al., 2018**). One-week post incubation, seedlings were randomly thinned out under a sterile bench to keep three plants per FlowPot. Plants were harvested at the vegetative stage five-weeks after seed sowing for all experiments.

### Shoot traits measurements

Shoots of Individual plants were cut and their fresh weight was first measured first. Shoots were disposed onto white paper, sealed with polyester non-sterile transparent film (VWR, USA), and scanned (Perfection V600 Photo, Epson). Canopy size, petiole length, leaf length-to-width ratio, and leaf numbers were measured using ImageJ (Fiji) (**Schindelin et al., 2012**).

### Microbial community profiling

For community profiling, plant roots (pool of three root systems per FlowPot) were thoroughly washed with sterile Milli-Q water and dried out with sterilized Whatman glass microfiber filters (GE Healthcare Life Sciences, USA). Matrix samples, corresponding to peat substrate in the pot without plant roots were also harvested (around 0.5 ml volume of peat per FlowPot). Plant roots and matrix samples were then transferred into individual lysing matrix E tubes (MP Biomedicals, Germany), frozen in liquid nitrogen and stored at −80°C. Samples were crushed using the Precellys®24 tissue lyzer at 6,200 rpm for 30 s (Bertin Technologies, Montigny-le-Bretonneux, France) and DNA isolation was performed using the FastDNA® SPIN for soil kit (MP Biomedicals, USA), as previously described (**Durán et al. 2018**). DNA concentration was quantified using the Quant-iT PicoGreen dsDNA assay kit (Invitrogen, Germany) and the fluorescence of dsDNA was measured by quantitative PCR (qPCR, IQ5 real-time PCR Thermocycler, Biorad, Munich, Germany) using the following parameters: 30 s at 25°C, 3×30 seconds at 25°C for measuring fluorescence, 30 seconds at 15 °C). Sample concentration was adjusted to 3.5 ng/μl with sterile water. Library preparation of individual samples involved a two-step PCR protocol. In the first PCR step (20 cycles, see **Durán et al. 2018**), the 25 μl reaction mix contained 3 μl sample DNA, 1x incomplete reaction buffer, 1.25 U DFS-Taq DNA Polymerase, 2 mM MgCl2, 0.3% BSA, 200 μM of dNTPs and 400 nM of each primer. Universal primers targeting the bacterial 16S rRNA V5-V6 region (799F/1192R), the fungal ITS1 (ITS1F/ITS2), and the oomycetal ITS1 (ITS1-O/5.8s-O-Rev) were used, as previously described (**Durán et al. 2018**). After the first PCR, enzymes and ssDNA were digested using a mixture of 1 μl Antarctic phosphatase, 1 μl Exonuclease I and 2.44 μl Antarctic phosphatase buffer that was added to 20 μl of the first-step PCR product (37 °C for 30 minutes and 85 °C for 15 minutes). After centrifugation, 3 μl of this solution was used as a template for a second PCR (10 cycles, see **Durán et al. 2018**). This step involved the same aforementioned PCR mix except that the universal primers were barcoded (reverse primers only) and included P5 and P7 Illumina adaptors. The full list of primers is available in **Durán et al. 2018.** After PCR, 5 μl PCR product for each barcoded sample was mixed with 5 μl 6×Orange G DNA Loading Dye (Orange G, 6X, Sigma, Germany) and run on a 1 % agarose gel in TAE 1X buffer. After control gel checking, the remaining PCR product (around 70 μl) was mixted with 20 μl Gel Loading Dye (Orange G) and run on a 1.5 % agarose gel for 2 hr at 80 V in order to excise the bacterial 16S rRNA band (500 bp). Bands were cut and purified using the QIAquick gel extraction kit (QIAGEN, Hilden, Germany) and DNA concentration was measured using the PicoGreen method as described above. For fungi and oomycetes, DNA purification was performed using the Agencourt AMPure XP-PCR Purification method (Beckman Coulter, Germany). DNA sample concentration was then quantified by the Quant-iT PicoGreen dsDNA method, as described above. For each microbial group (bacteria, fungi and oomycetes), 30 ng DNA of each of the barcoded samples were mixed, resulting in three pooled samples that were purified twice (AMPure XP-PCR Purification, Beckman Coulter, Germany). The final DNA concentration of each pool was measured using the Quantus™ Fluorometer (Promega, Germany) and an equal quantity of DNA (i.e., 100 ng) was used to assemble a single library from the three bacterial, fungal, and oomycetal DNA pools. Finally, paired-end Illumina DNA sequencing was performed using the Illumina MiSeq system available on site at the Max Planck Institute for Plant Breeding Research.

### 16S rRNA and ITS read processing

Paired amplicon sequencing reads were joined using Qiime (join_paired_reads, **Caporaso et al., 2010**). In case of ITS reads, for un-joined read pairs, corresponding forward reads were retained for demultiplexing. Demultiplexing and quality filtering was done using Qiime (split_libraries_fastq, phred=30). All quality filtered and demultiplexed reads were trimmed to an equal length. Reference sequences were obtained from all used strains (bacteria = 183, fungi=24, oomycetes=7, see Table S1) using the respective resources of sequenced genomes, if available. Reference sequences were then trimmed to contain only the amplified region. Trimmed reference sequences that were 100% identical, were grouped in so called strain variants. This led to 115 bacterial, 24 fungal and 7 oomycetal strain variant sequences. Reference sequences of strain variants were mapped against all trimmed amplicon reads using usearch (**Edgar et al., 2010**), allowing for one mismatch (usearch-usearch_global, with max diff = 1). Unmapped reads were discarded. Count tables were made from these mapping results.

### Microbial community profile statistical analysis

To calculate alpha diversity indices, count tables were rarefied to 1000 reads. Significant differences between alpha diversity indices were determined by an ANOVA followed by *post hoc* Tukey’s HSD (*P* < 0.05). Distance matrices were calculated by using a normalized count table (CSS, **Paulson et al., 2013**), which was used for Bray Curtis distance calculation. Distance tables were used as an input for principal coordinate analysis (PCoA) and constrained analysis of principal coordinates (CAP, capscale function in R). The condition function was used to correct for batch effects and when applicable for treatments (LP/NC) when performing CAP. Significance of CAP results was tested using ANOVA (anova.cca in R, *P* < 0.05). Partial least square discriminant analysis (PLS-DA) were used to discriminate groups of samples based on the composition of their microbial community. The PLS-DA consists of a partial least squares (PLS) regression analysis where the response variable is categorical (y-block; describing the grouping factor), expressing the class membership of the statistical units (**Sjöström et al., 1986, Sabatier et al., 2003**). Raw OTU tables were first scaled before using the cppls function in the package “pls”. The PLS-DA procedure includes a cross-validation step producing a p value that expresses the validity of the PLS-DA method regarding the data set (function MVA.test in package “RVAideMemoire”). The PLS-DA procedure also expresses the statistical sensitivity indicating the modeling efficiency in the form of the percentage of misclassification of samples in categories accepted by the class model (function MVA.cmv in package “RVAideMemoire”). To better plot differences in abundances for individual strains across different sets of experiments, relative abundances were rescaled to be in a range from 0 to 1. For this read counts per sample were transformed to relative abundances by division of total sum of reads per sample. Then for all samples, belonging to one experiment, RA of each individual strain variant across desired samples to analyze (e.g. only root samples, only wt and mutant samples) were rescaled using the general equation x-(min(x))/(max(x)-min(x)), where x represents all RA values per strain variant from one experiment. Strain variants that where not consistently found across samples were discarded from this analysis. Enrichment of strain variants across LP and NC conditions was calculated using the following steps (all functions from R package “EdgeR”). Raw read counts were normalized using TMM normalization (“calcNormFactors”). A generalized linear model was fitted to integrate the batch effect (“glmFit”). Enrichment was then determined with a likelihood ratio test (“glmLRT”, *P* < 0.05). A predictive Support Vector Machines model with a linear kernel was trained (*SVC* function of Scikit-learn, **Pedregosa et al. 2011**) to link standardized relative abundances of bacteria to plant rescue. Recursive Feature Elimination with Cross Validation (*RFECV* function) was performed to identify the smallest set of bacteria discriminating the two plant phenotypes, and to estimate the model accuracy using a leave-one-out approach (K-Folds cross-validator, with K equaling the number of samples). The species identified in the RFECV were then used to compute a PLS-DA.

### Transcriptome sequencing experiments

Shoot and roots of five-weeks-old *A. thaliana* Col-0, *myc2-3* and *jin1-8* pMYC2:MYC2-FLAG plants growing in the FlowPot system were harvested separately at 10 a.m. Roots were washed quickly (< 1 min) to detach the surrounding peat matrix using 10% RNAlater (QIAGEN, Valencia, CA) in 1X PBS as the capture buffer to mitigate RNA degradation. Total RNA from all samples was extracted using the RNeasy® Plant Mini Kit (QIAGEN, Germany). DNA removal was performed using RNase-Free DNase Set (QIAGEN, Germany). Eleven samples from *myc2-3* plants detected more than 50% reads mapped to latent virus genome. Based on the hierarchical relationship between samples from *myc2-3* mutant (**Figure S6D**), the contamination of latent virus did not significantly affect plant transcriptome. RNA-Seq libraries were prepared and sequenced at the Max Planck Genome Centre (Cologne, Germany) with an Illumina HiSeq2500. For samples from Col-0 plants, run conditions were 1×150 bp (single read) and total reads per sample were 20 million. For samples from *myc2-3* plants, run conditions were 1×150 bp (single read) and total reads per sample were 55 million. For samples from *jin1-8* pMYC2:MYC2-FLAG plants, run conditions were 1 ×150 bp (single read) and total reads per sample were 10 million.

### Transcriptome sequencing data analysis

The FastQC suite was performed to check the quality of the sequenced reads (http://www.bioinformatics.babraham.ac.uk/projects/fastqc/). Subsequently, the RNA-Seq reads were mapped to the annotated genome of *A. thaliana* (TAIR10) using Tophat2 (**Kim et al., 2013**, tophat2 -p 20 -a 10 -g 10) with Bowtie2 (**Langmead et al., 2012**) building genome index. The mapped RNA-Seq reads were subsequently transformed into a fragment count per gene per sample using the htseq-count script (s=reverse, t=exon) in the package HTSeq (**Anders et al., 2015**). Count tables of Col-0 samples were concatenated to one count matrix. Count tables of *myc2-3* and *jin1-8* pMYC2:MYC2-FLAG samples were concatenated to one count matrix. To normalize count data, raw counts were first calculated normalization factor via TMM-normalization. Then the genes with more than 100 counts were extracted. The extracted TMM-normalized count data were transformed to log2cpm via the voom function in the limma package (**Ritchie et al., 2015**) in R (R version 3.6.3). Subsequently, log2cpm data were used to calculate log2 fold changes and p-values (F-test) for individual comparisons. Resulting p-values were adjusted for false discoveries (FDR) due to multiple hypotheses testing via the Benjamini-Hochberg procedure. To identify significant different expression genes, a threshold of |log2FC| ≥ 1 and FDR < 0.05 was applied. Heatmaps of significantly regulated genes expression profiles were generated using the pheatmap package (**Kolde 2015**) in R. The Euclidean distance was used to show the distance among clustering rows. The values in heatmaps were scaled by row. Gene Ontology (GO) enrichment was conducted using the enrichGO function in the clusterProfiler package (**Yu et al., 2012**) in R. Biological Process (BP) and 0.05 p-values cutoffs were chosen. P-values were adjusted via the Benjamini-Hochberg method.

### Variance partitioning of microbial community composition and transcriptomic profiles

Variance partitioning between experimental factors was tested with a permutational ANOVA approach with the Adonis function (R package vegan) in all experiments.For the stress experiment, the effect of compartment and light condition factors on microbial community composition was tested in a global model. The effect of light conditions was also tested in separated models for roots and shoots samples. These models where constructed separately on each of the bacterial, fungal and oomycetal datasets using Bray-Curtis dissimilarity matrices between pairs of samples produced with the vegdist function (R package vegan). For the mutant experiment, the effect of genotype, compartment and light condition factors on bacterial community composition was tested in a global model. The effect of genotype and light conditions was also tested in separated models of roots and matrix samples. These models where constructed using Bray-Curtis dissimilarity matrices between pairs of samples produced with the vegdist function (R package vegan). For the transcriptomic experiments, the effect of experimental factors on the transcriptomic expression profile of *A. thaliana* plants was tested using Euclidean distance matrices between pairs of samples produced with the vegdist function (R package vegan). Models were constructed separately for Col-0 and *myc2-3/ jin1-8* pMYC2:MYC2-FLAG plants as these were harvested in separated experiments. For Col-0 plants the effect of compartment, microbes’ inoculation and light conditions on the plant transcriptomic profile was tested in a global model. The effect of microbes’ inoculation and light conditions was also tested in separated models for roots and shoots samples. For *myc2-3/ jin1-8* pMYC2:MYC2-FLAG plants, the effect of compartment, genotype, microbe inoculation and light conditions on the plant transcriptomic profile was tested in a global model. The effect of genotype, microbes and light was also tested in separated models for roots and shoots samples. These models were also constructed for *myc2-3* and *jin1-8* pMYC2:MYC2-FLAG datasets separately to test the effect of compartment, microbes’ inoculation and light effect independently of the genotype effect.

### Pathogen inoculation and symptom quantification

For *Botrytis cinerea* B05.10 (*Bc*) inoculation of *A. thaliana*, the spores were diluted in Vogelbuffer (in 1 L: 15 g of Suc, 3 g of Na-citrate, 5 g of K_2_HPO_4_, 0.2 g of MgSO_4_·7H_2_O, 0.1 g of CaCl_2_·2H_2_O, and 2 g of NH_4_NO_3_) to 5 × 10^5^ spores ml^-1^. For droplet inoculations, 2 μl droplets containing 1×10^3^ spores were applied to each leaf of four-week-old *A. thaliana* grown in the presence/absence of the BFO SynCom under either NC or LP. The entire infection processes were conducted under a sterile clean bench. After five days post pathogen inoculation in the FlowPot system, shoots were cut, washed using Milli-Q water twice, and dried out with sterilized Whatman glass microfiber filters (GE Healthcare Life Sciences). Shoots were weighed, placed into 2 ml sterile Eppendorf tubes, and snap-frozen in liquid nitrogen. For quantification of fungal growth, DNA from plant shoots was extracted using the DNeasy® Plant Mini Kit (QIAGEN, Germany). The relative amounts of *Bc* and *A. thaliana* DNA were determined by qPCR, employing specific primers for cutinase A and SKII, respectively, as described previously (Liu et al. 2017). *Pseudomonas syringae pv. tomatao* DC3000 (*Pst*) inoculation was carried out by spaying *Pst* at 0.2 OD in 10 mM MgCl_2_ on *A. thaliana* leaves of four weeks-old *A. thaliana* grown in the presence/absence of the BFO SynCom under either NC or LP. The entire infection process was conducted under a sterile clean bench. After five days post pathogen inoculation in the FlowPot system, plant shoots were cut, washed using 70% EtOH once and rinsed in Milli-Q water twice. Plant shoots were dried with sterilized Whatman glass microfiber filters and weighted under a sterile clean bench. Then plant shoots were put into 2 ml sterile Eppendorf tubes containing 1.5 ml 10 mM MgCl_2_/0.01% Silwet. The tubes were then directly shaken at 650 rpm for 1 h at 28°C. For colony counting, a series of dilutions were conducted with 10 mM Mgcl_2_ to 10^-1^,10^-2^,10^-3^,10^-4^, and 10^-5^. Then, 20 μl of diluted liquid was spotted on NYGA plates, counting colonies after two days incubation at 28 °C.

### Immunoblot analysis

To analyze MYC2 protein levels of plants under NC/ LP in the presence or absence of BFO, the shoots and roots of *A. thaliana* col-0 and *jin1-8* pMYC2:MYC2-FLAG plants, grown in the FlowPot system under NC/ LP conditions as described above, were harvested separately into 2 ml Eppendorf tubes with one 3.2 mm stainless-steel beadafter 5 weeks-BFO/ +BFO inoculation and frozen in liquid nitrogen. Samples were ground to powder in liquid nitrogen, and boiled in 150 μl 2xLaemmli buffer for 10 min at 95°C. The proteins were resolved on 10% SDS-PAGE (1610156, Bio-Rad) and transferred using the wet transfer method onto a nitrocellulose membrane (10600001, GE Healthcare Life Sciences). Monoclonal Anti-β-Actin-Peroxidase antibody produced in mouse (A3854, Sigma-Aldrich, Germany) was used to detect Actin, and to adjust the total protein concentration. Monoclonal ANTI-FLAG® M2-Peroxidase antibody produced in mouse (A8592, Sigma-Aldrich, Germany) was used to detect MYC2 protein. Both antibodies were used at a dilution of 1:5,000 (1xTBST, 5% milk powder). Detection of the signal was performed with SuperSignal West Pico and Femto (34080 and 34095, ThermoFisher Scientific, Germnay) using ChemiDoc Imaging Systems (Bio-Rad,Germany). The protein concentration was calculated based on the band thickness using Fiji.

### Phytohormone measurement

To measure phytohormone of plants under NC/ LP in the presence or absence of BFO, the shoots and roots of *A. thaliana* col-0 and *myc2-3* plants, grown in the FlowPot system under NC/ LP conditions as described above, were harvested separately into 2 ml Eppendorf tubes with one 3.2 mm stainless-steel bead. Phytohormone concentrations were determined from 50 mg of tissue fresh weight. Sample processing, data acquisition, instrumental setup, and quantifications (using 5 ng [^2^H]_6_ JA, 0.75 ng [^2^H]_2_-JA-Ile, 30 ng [^2^H]_5_-OPDA, and 1.5 ng [^2^H]_4_-SA as internal standards per sample) were performed as described (**Ziegler et al., 2014**, **Liu et al., 2015**).

## QUANTIFICATION AND STATISTICAL ANALYSIS

Data collection and analysis was performed under blinded conditions in all experiments. All statistical analyses were performed using the R environment (R version 3.6.3). *P* < 0.05 were considered as significant. Individual statistical tests used for each figure are available at https://github.com/ShijiHou/Light-limitation-Paper.

## DATA AVAILABILITY

Sequencing reads from microbiota reconstitution experiments (MiSeq 16S rRNA and ITS reads) have been deposited in European Nucleotide Archive (ENA, PRJEB40980 – bacteria, PRJEB40981 – fungi, PRJEB40982 – oomycetes). Sequencing reads from transcriptome sequencing experiments have been deposited in Gene Expression Omnibus (GEO, GSE160106 – Col-0 data, GSE160115 – *myc2-3* and *jin1-8* pMYC2:MYC2-FLAG data). All scripts for computational analysis and corresponding raw data are available at https://github.com/ShijiHou/Light-limitation-Paper.

## SUPPLEMENTARY FIGURES

**Figure S1.**
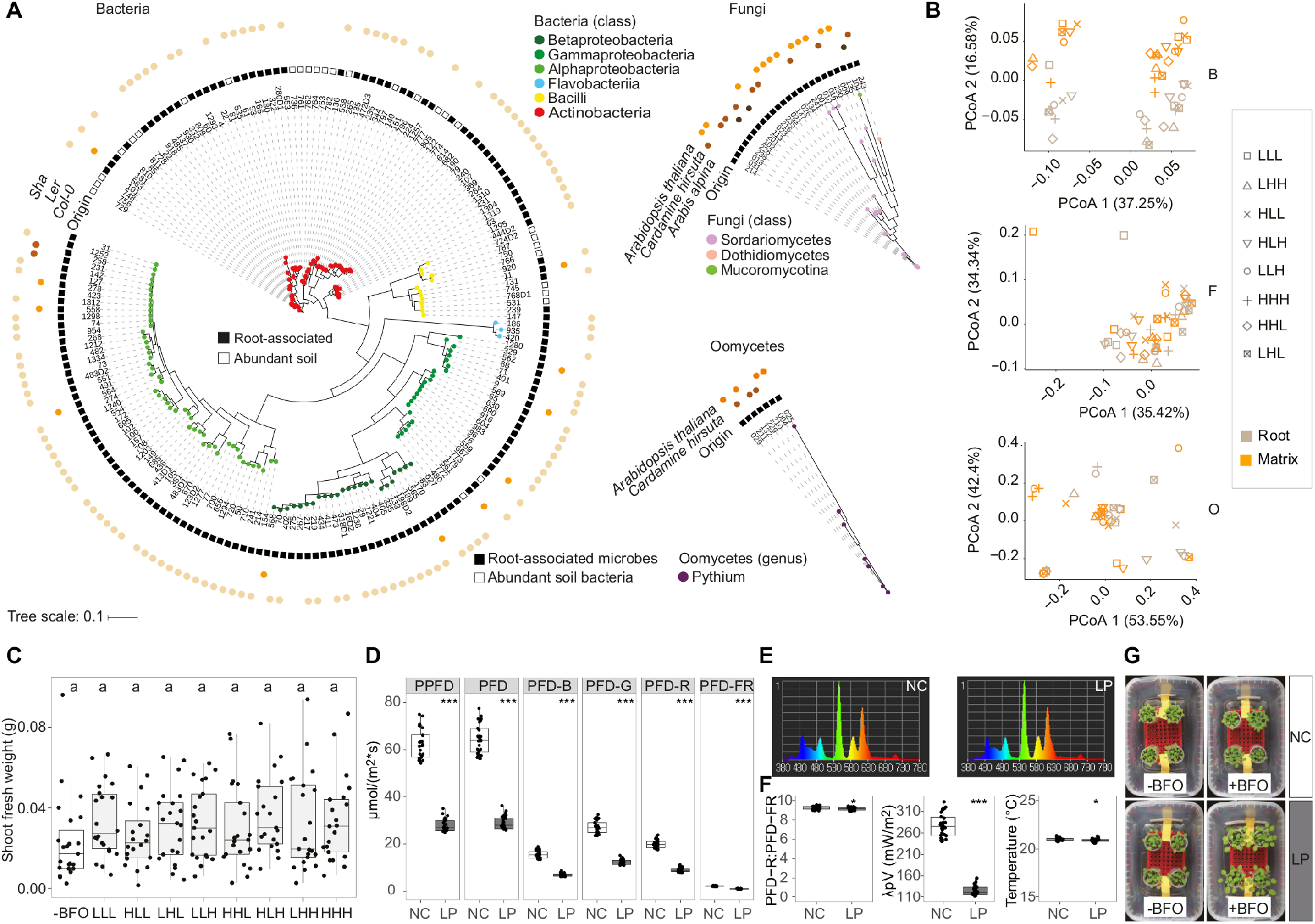
BFO SynCom composition, stability, and effect on plant growth. **(A)** 183 Bacterial, 24 fungal, and 7 oomycetal (BFO) strains were used to reconstitute a synthetic *A. thaliana* root microbiota. Taxonomic trees were constructed based on bacterial 16s rRNA V5V7 sequences and fungal/oomycetal ITS1 sequences. The trees were edited in iTol (**Letunic and Bork 2019**) and tracks represent (from the outside to the inside): name of the host plant/genotype (Shakdara: Sha, Landsberg erecta: Ler and Columbia-0: Col-0 are different *A. thaliana* accessions), origin of isolation (soil or root compartment), and microbial strain identification number. Circles at the extremity of the branches are colour coded according to strain taxonomic assignment at the class level for B and F, and at the genus level for O. **(B)** Impact of the relative abundance (RA) of B, F, and O on BFO community composition in roots and surrounding peat matrix in the FlowPot system. Data were visualized using a principal-component analysis (PcoA) and the first two dimensions of the PCoA were plotted based on Bray-Curtis dissimilarities. Samples are colour-coded according to the compartment, and BFO ratio conditions are depicted with different symbols. For inoculation, B, F, and O communities were assembled separately and microbial suspensions were adjusted to OD = 0.5 (2 ml/50 ml) for B and to 100 mg/50 ml for F and 40 mg/50 ml for O (high RA, H), or to OD = 0.5 (0.2 ml/50 ml) for B and to 10 mg/50 ml for F and 4 mg/50 ml for O (i.e., 10 times dilution, low RA, L). Eight possible combinations of B-to-F-to-O RA ratios were then assembled and used to repopulate sterile peat in the FlowPot system. Plant roots and surrounding peat matrix were harvested 5 weeks post BFO inoculation. Note that the BFO RA ratio B:L, F:L, O:L (LLL) was selected and used for all following experiments. Three independent biological replicates (n = 51 samples). **(C)** Shoot fresh weight (in g) of five-week-old *A. thaliana* inoculated with BFO at mixed at different B-to-F-to-O RA ratios or without BFO (-BFO). Three independent biological replicates (n = 191 plants). Letters indicate statistical significance corresponding to Kruskal-Wallis with Dunn’s *post hoc* test (α = 0.05). **(D)** Light parameters measured in the FlowPot system at different positions in the plant growth chamber (λp = 545 nm). PPFD: Photosynthetic Photon Flux Density defined in 400~700nm, PFD: Photosynthetic Photon Flux Density defined in 380~780nm, PFD-B: Photosynthetic Photon Flux Density in blue field (400~500nm), PFD-G: Photosynthetic Photon Flux Density in green field (500~600nm), PFD-R: Photosynthetic Photon Flux Density in red field (600~700nm), PFD-FR: Photosynthetic Photon Flux Density in far-red field (700~780nm). Asterisks indicate statistical significance (Mann-Whitney U test, *** *P* < 0.001). **(E)** Spectrum in the FlowPot system under NC (normal light, left) and LP (low PAR, right). **(F)** R:FR ratio (left), λpV (Peak wavelength value: the highest power in the measured spectrum, middle), and temperature (right) measured in the FlowPot system at different positions in the plant growth chamber under different light conditions (NC: normal light and LP: low PAR). Asterisks indicate statistical significance (T-test for R:FR ratio, Mann-Whitney U test for Peak wavelength value and temperature, *** *P* < 0.001, * *P* < 0.05). **(G)** Representative pictures showing five-week-old plants growing in the FlowPot system under different light (NC: normal light, LP: low PAR) and BFO (-BFO, +BFO) conditions.

**Figure S2.**
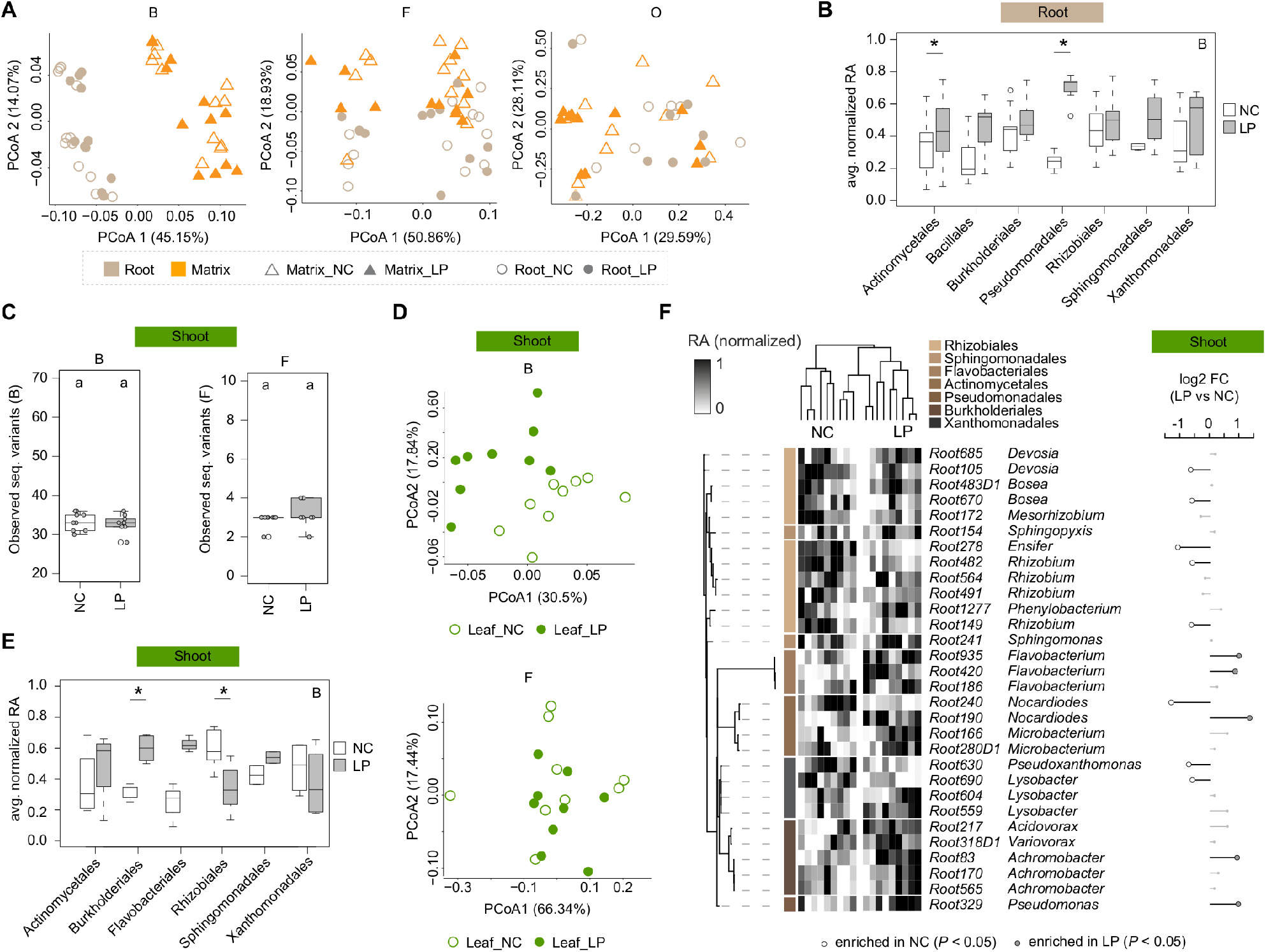
Influence of compartment and light conditions on BFO community composition. **(A)** Effect of “Compartment” and “Light” on B, F, and O community composition in roots and surrounding peat matrix in the FlowPot system. Data were visualized using a principal-component analysis (PCoA) and the first two dimensions of the PCoA were plotted based on Bray-Curtis dissimilarities. Samples were depicted according to the compartment (matrix: orange triangles; root: brown circles) and the light conditions (normal light condition, NC, open symbols; Low PAR, LP, filled symbols). For inoculation, B, F, and O communities were assembled separately and microbial suspensions were adjusted to OD= 0.5 for B, and to 0.2 mg/ml for F, and to 0.08 mg/ml for O, corresponding to the BFO ratio LLL defined in **Figure S1**. Three independent biological replicates (B: n = 47 samples, F: n = 48, O: n = 37). **(B)** Mean normalized RA in LP/NC for bacterial strain variants sorted by taxonomic order. Only strain variants that were consistently found across all samples (n=24) were selected. Bacterial orders with less than one member were not considered (strain variants: Act. n = 28, Bac. n= 6, Bur. n= 9, Pse. n= 7, Rhi. n= 20, Sph. n= 4, Xan. n= 8). Significant differences between norm. RA in LP and NC are shown (Mann-Whitney-U test, *P* < 0.05). **(C)** Number of bacterial (B, left) and fungal (F, right) strain variants detected in plant shoots. Three independent biological replicates (n = 18 samples). Letters indicate statistical significance corresponding to Tukey’s HSD (α = 0.05). NC: normal light condition, LP: low photosynthetically active radiation. **(D)** Effect of “Compartment” and “Light” on B and F community composition in shoots in the FlowPot system visualized by PCoA (Upper panel: B, Lower panel: F, Empty circles = NC, filled circles LP). Three independent biological replicates (B: n = 18 samples, F: n = 18)**. (E)** Mean normalized RA in LP/NC for bacterial strain variants detected in plant shoots sorted by taxonomic order. Only strain variants that were consistently found across all samples (n=18) were selected. Bacterial orders with less than one member were not considered (strain variants: Act. n = 4, Bur. n= 5, Fla. n= 3, Rhi. n= 10, Sph. n= 2, Xan. n= 4). Significant differences between norm. RA in LP and NC are shown (Mann-Whitney-U test, *P* < 0.05). **(F)** Normalized RA of prevalent B strain variants detected in shoots across NC and LP conditions. Sample-wise normalized RA was calculated for each strain variant and is depicted in the heatmap next to the taxonomic tree. Only strain variants with an average RA ≥ 0.1% across samples were considered (n = 18 samples). The taxonomic tree was constructed based on a maximum likelihood method using full length 16S rRNA sequences. Sample-wise log2 fold-change in RA measured for each prevalent strain variant between LP and NC in shoot samples. Differential RA with statistically significant *P*-values are shown (edgeR, GLM, *P* < 0.05).

**Figure S3.**
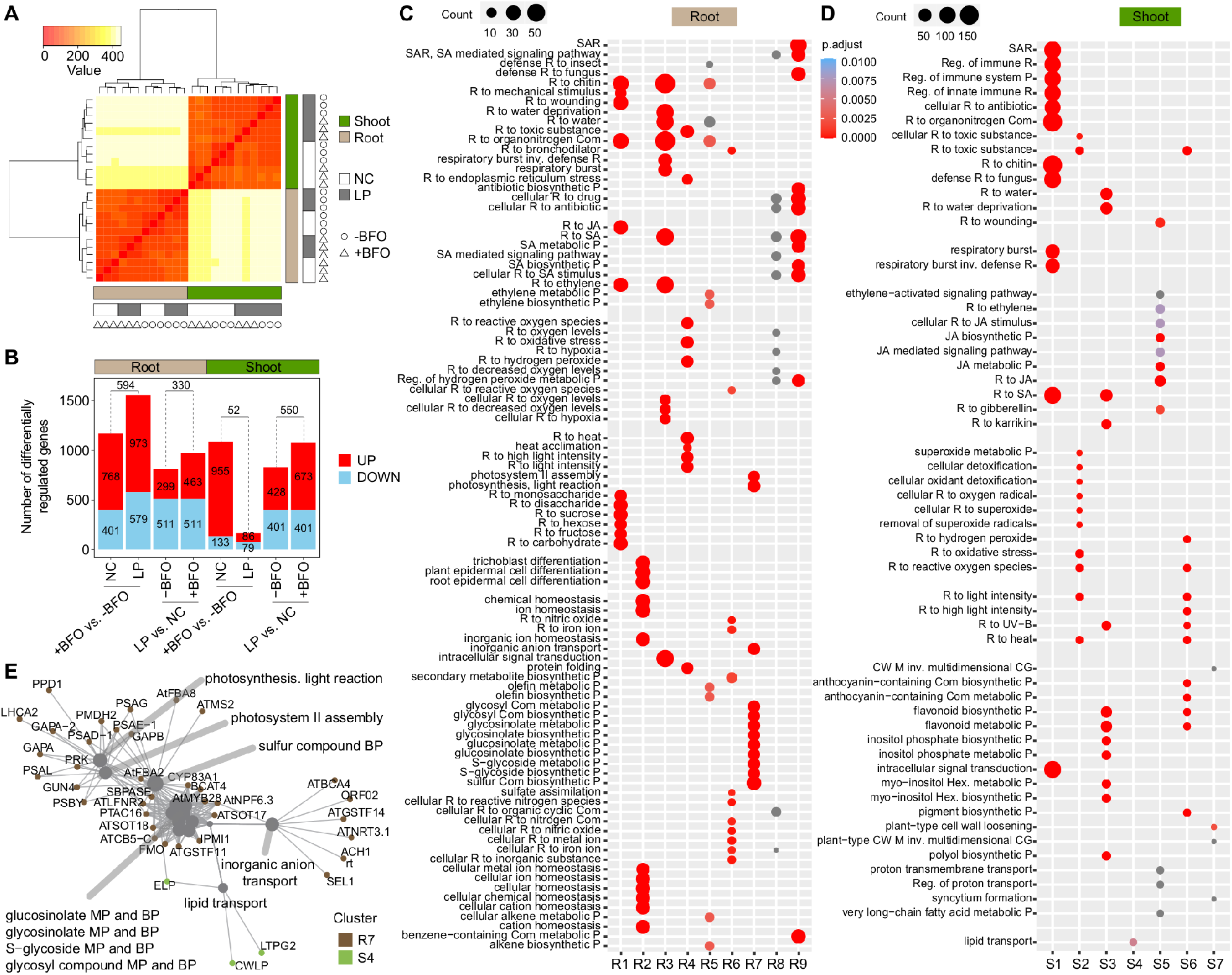
Transcriptional reprogramming in shoot and root in response to light and BFO. **(A)** Heatmap showing the hierarchical relationship between samples used for RNAseq. Three independent biological replicates (n = 24 samples). NC: normal light condition, LP: low PAR. -BFO: no microbes (i.e., germ-free), +BFO: with microbes. B: bacteria, F: fungi, O: oomycetes. The colour key on the top left side stands for sample distances (using *“dist”* function in “DESeq” R package to the transpose of the transformed count matrix to get sample-to-sample distances) corresponding to the colour range in the heatmap. **(B)** Bar plot showing the numbers of differentially regulated genes among all possible pairwise comparisons. Red colour and blue colour filled bars represent significantly up- and down-regulated genes between the corresponding comparisons, respectively. The numbers highlighted inside the bars indicate the total number of up-or down-regulated genes between the corresponding comparisons. The numbers above the bars represent the numbers of genes shared between the corresponding conditions. **(C)** and **(D)** Gene Ontology (GO) term enrichment depicting the top 12 most significantly enriched GO terms (Hypergeometric test with Bonferroni correction, *P* < 0.05) detected in all clusters for root samples (D) and shoot samples (E). The size of point reflects the amount of gene numbers enriched in this GO term. The colour of point means the p value ((Hypergeometric test with Bonferroni correction). R: response, Reg. Regulation, ET: ethylene, SA: salicylic acid, JA: jasmonic acid, SAR: systemic acquired resistance, CW: cell wall, CG: cell growth, P: process, Com: compound, Hex: hexakisphosphate, inv.: involved in. **(E)** Gene-concept network (cnetplot function in R) depicting linkages between genes and associated top 12 most significantly enriched GO terms detected in clusters R7 (root) and S4 (shoot). Each node represents a gene and is colour-coded according the different cluster names. MP: metabolic process. BP: biosynthetic process.

**Figure S4.**
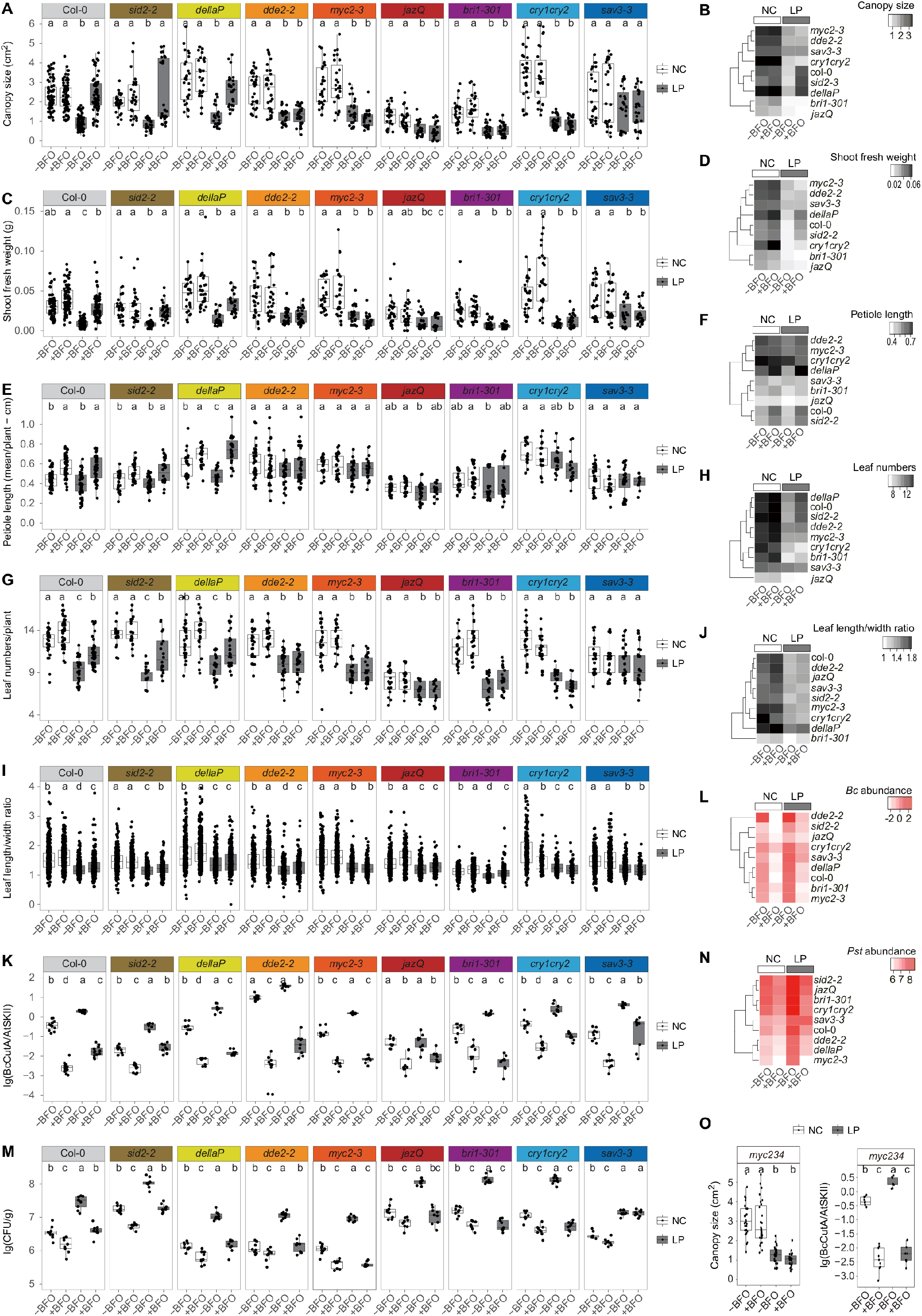
BFO-induced growth over defence in LP depends on JA and BR pathways. **(A)** Canopy size (in cm^2^) of *A. thaliana* Col-0 and eight mutants grown under NC and LP, in the absence (-BFO) or presence of the BFO SynCom (+BFO), in the FlowPot system. The canopy size of five-weeks old plants was measured using Fiji and data from three independent biological replicates are shown (n = 1188 plants across replicates and genotypes). Statistical significance across conditions for each genotype is indicated with letters (Kruskal-Wallis with Dunn’s *post hoc* test, α = 0.05). NC: normal light condition, LP: low PAR. **(B)** Summary heatmap depicting the mean canopy size (in cm^2^) measured across plant genotypes and conditions. **(C)** Shoot fresh weight (in g) of five-week-old *A. thaliana* Col-0 and eight mutants grown under NC and LP, in the absence (-BFO) or presence of the BFO SynCom (+BFO), in the FlowPot system. Three independent biological replicates are shown (n = 1188 plants across replicates and genotypes). Statistical significance across conditions for each genotype is indicated with letters (Kruskal-Wallis with Dunn’s *post hoc* test, α = 0.05). **(D)** Summary heatmap depicting the mean shoot fresh weight (in g) measured across plant genotypes and conditions. **(E)** Mean petiole length (in cm) of five-week-old *A. thaliana* Col-0 and eight mutants grown under NC and LP in the absence (-BFO) or presence of the BFO SynCom (+BFO), in the FlowPot system. Three independent biological replicates are shown (n = 1095 plants across replicates and genotypes). Statistical significance across conditions for each genotype is indicated with letters (ANOVA followed by *post hoc* Tukey’s HSD). **(F)** Summary heatmap depicting the mean value of mean petiole length per plant (in cm) measured across plant genotypes and conditions. **(G)** Total leaf number of five-week-old *A. thaliana* Col-0 and eight mutants grown under NC and LP, in the absence (-BFO) or presence of the BFO SynCom (+BFO), in the FlowPot system. Three independent biological replicates are shown (n = 925 plants across replicates and genotypes). Statistical significance across conditions for each genotype is indicated with letters (Kruskal-Wallis with Dunn’s *post hoc* test, α = 0.05). **(H)** Summary heatmap depicting the mean total leaf number measured across plant genotypes and conditions. **(I)** Leaf length/width ratio of five-week-old *A. thaliana* Col-0 and eight mutants grown under NC and LP, in the absence (-BFO) or presence of the BFO SynCom (+BFO), in the FlowPot system. Three independent biological replicates are shown (n = 9,708 leaves across replicates and genotypes). Statistical significance across conditions for each genotype is indicated with letters (Kruskal-Wallis with Dunn’s *post hoc* test, α = 0.05). **(J)** Summary heatmap depicting the mean value of mean leaf length/width ratio measured across plant genotypes and conditions. **(K)** *B. cinerea* B05.10 (*Bc*) growth in leaves of *A. thaliana* Col-0 and eight mutants grown under NC and LP, in the absence (-BFO) or presence of the BFO SynCom (+BFO), in the FlowPot system. *Bc* growth was quantified by qPCR in *A. thaliana* leaves five days post pathogen inoculation. For inoculation, 2 μl droplets containing 1×10^3^ spores were applied to leaves of four weeks-old *A. thaliana* grown in the presence/absence of the BFO SynCom under either NC or LP. The relative growth of *Bc* and *A. thaliana* was determined by amplification of the *BcCutinaseA* gene and the *AtSkII* genes, respectively. Three independent biological replicates are shown (n = 336 samples across replicates and genotypes). Statistical significance across conditions for each genotype is indicated with letters, (ANOVA followed by *post hoc* Tukey’s HSD, α = 0.05). **(L)** Summary heatmap depicting relative *Bc* growth (lg(BcCutA/AtSKII)) measured across plant genotypes and conditions. **(M)** *Pseudomonas syringae pv. tomato* DC3000 (*Pst)* growth in leaves of *A. thaliana* Col-0 and eight mutants grown under NC and LP, in the absence (-BFO) or presence of the BFO SynCom (+BFO), in the FlowPot system. *Pst* growth was quantified by colony counting in *A. thaliana* leaves five days post pathogen inoculation. *Pst* inoculation was carried out by spaying *Pst* at OD 0.2 in 10 mM MgCl2 on leaves of four weeks-old *A. thaliana;* grown in the presence/absence of the BFO SynCom under either NC or LP. The data from three independent biological replicates are shown (n = 324 samples across replicates and genotypes). Statistical significance across conditions for each genotype is indicated with letters (ANOVA followed by *post hoc* Tukey’s HSD, α = 0.05). **(N)** Summary heatmap depicting relative *Pst* growth (lg(CFU/g)) measured across plant genotypes and conditions. **(O)** Canopy size (in cm^2^, left) and *B. cinerea* B05.10 (*Bc*) growth (right) in leaves of five-week-old *myc234* mutants grown NC and LP, in the absence (-BFO) or presence of the BFO SynCom (+BFO), in the FlowPot system. *Bc* growth was quantified by qPCR in *A. thaliana* leaves five days post pathogen inoculation. For inoculation, 2 μl droplets containing 1×10^3^ spores were applied to leaves of four weeks-old *A. thaliana* grown in the presence/absence of the BFO SynCom under either NC or LP. The relative growth of *Bc* and *A. thaliana* was determined by amplification of the *BcCutinaseA* gene and the *AtSkII*, respectively. Three independent biological replicates are shown (Canopy size: n = 108; *Bc* growth: n = 36). Statistical significance across conditions for each genotype is indicated with letters (Canopy size: Kruskal-Wallis with Dunn’s *post hoc* test, α = 0.05; *Bc* growth: ANOVA followed by *post hoc* Tukey’s HSD, α = 0.05).

**Figure S5.**
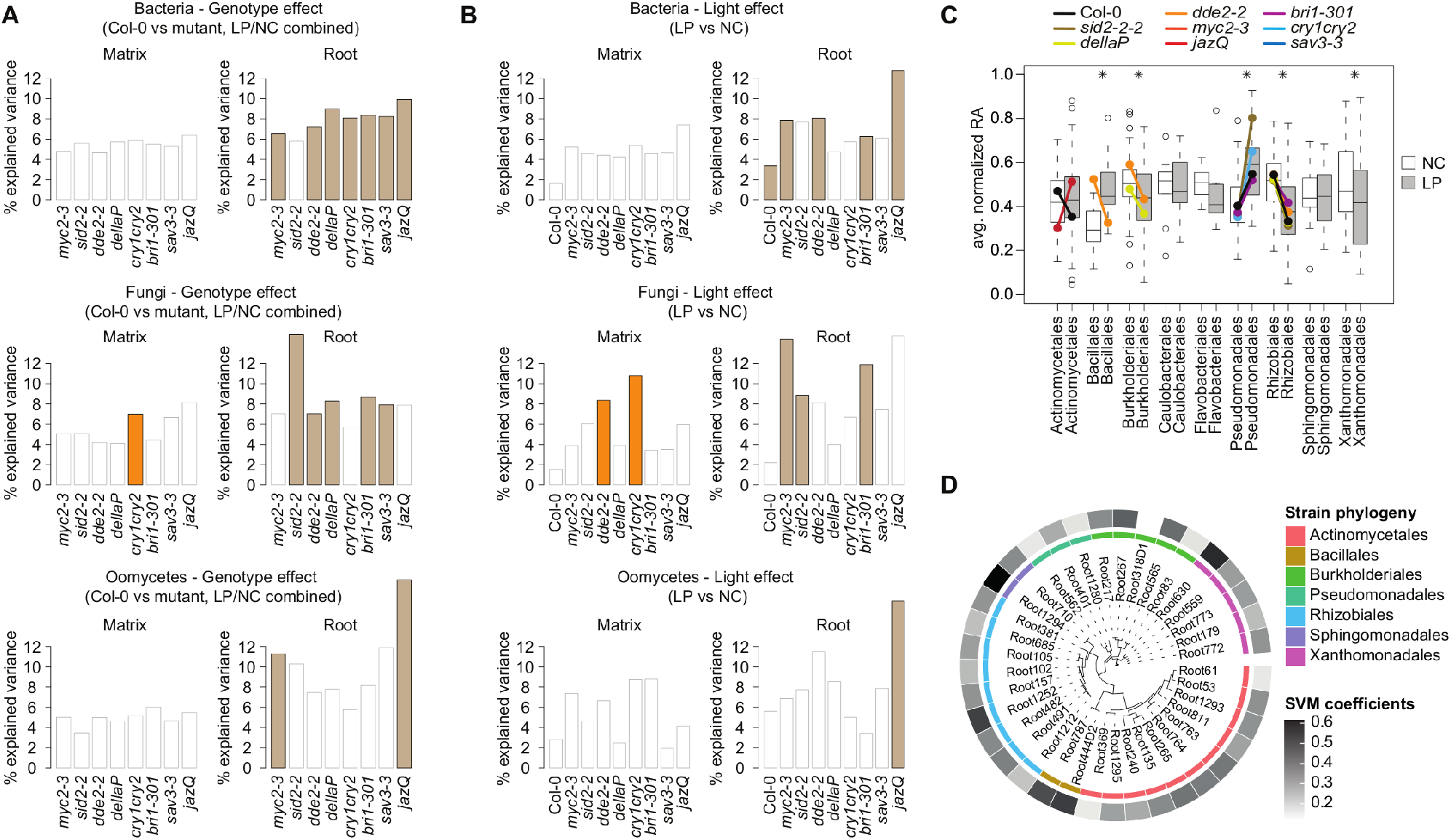
Genotype-dependent microbial community shifts between LP and NC. **(A)** Genotype effect for all datasets (Bacteria upper panel, Fungi middle panel, Oomycetes lower panel). Explained variance between genotype and Col-0 samples was calculated using CAP. Using all matrix (left) and root (right) samples respectively (including both LP/NC samples). Significance is highlighted with coloured boxes (ANOVA permutation test, *P* < 0.05). **(B)** Light effect for all datasets (Bacteria upper panel, Fungi middle panel, Oomycetes lower panel) across all genotypes. Explained variance between LP and NC samples was calculated using CAP. Using all matrix (left) and root (right) samples respectively. Significance is highlighted with coloured boxes (ANOVA permutation test, *P* < 0.05). **(C)** Mean normalized RA in LP/NC samples for bacterial strain variants for all genotypes, sorted by taxonomical order. Boxplot depicting all genotypes together, to show general trends. Coloured dots depict means within genotypes that are significantly different (Mann-Whitney-U test, *P* < 0.05). **(D)** 16S rRNA-based phylogeny of the 37 bacterial variants that discriminate best BFO-induced plant phenotypes under LP conditions (i.e., rescued vs. not rescued, SVM-RFE, R2=0.83). The outer circle displays the coefficients of the model vectors, reflecting the contribution of individual strains to the segregation of the two phenotypic groups.

**Figure S6.**
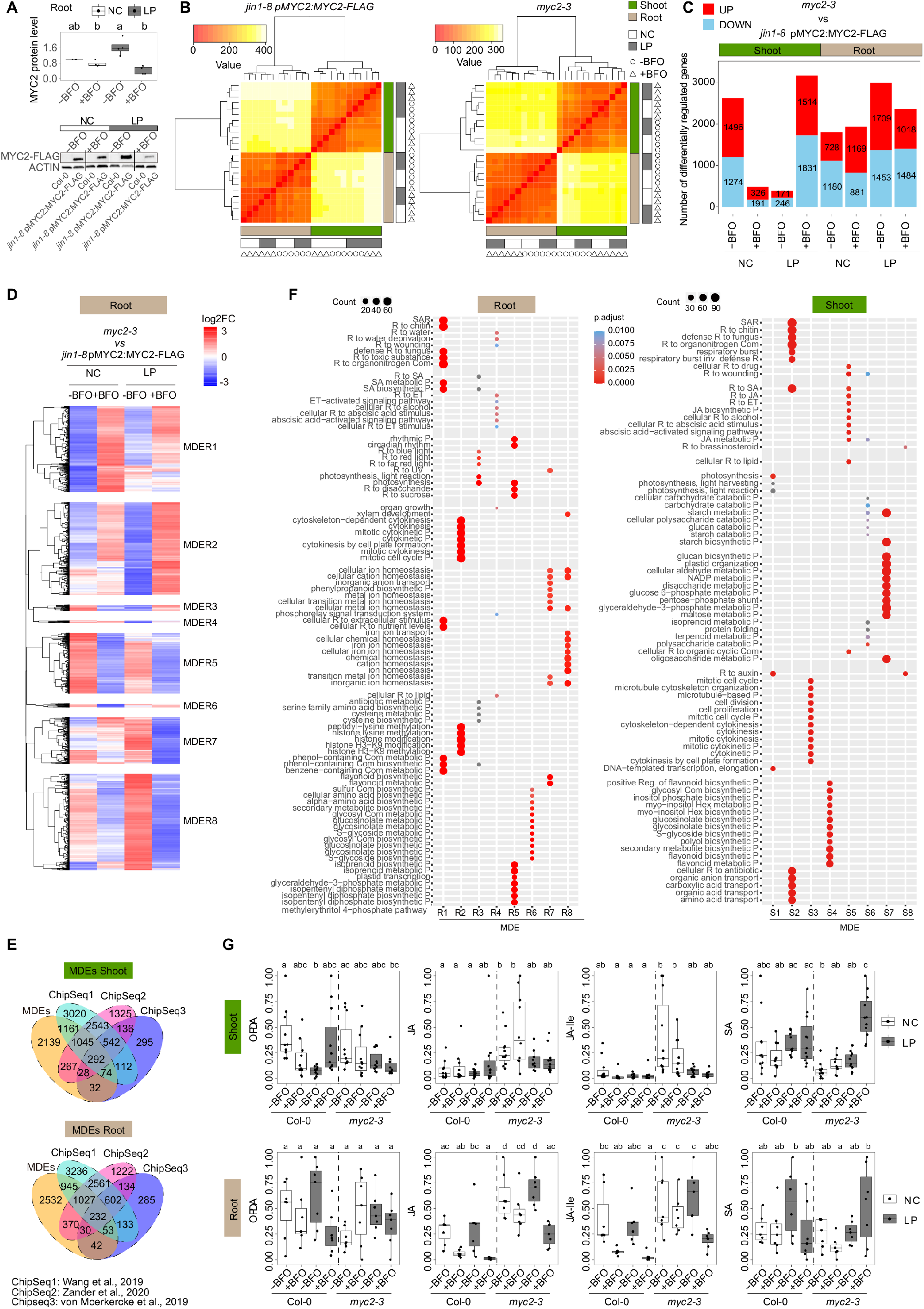
MYC2-dependent prioritization of BFO-induced growth over defence in LP. **(A)** Quantified MYC2 protein levels in roots of five-week-old *jin1-8* pMYC2:MYC2-FLAG line grown in the FlowPot system in the absence (-) or presence (+) of the BFO SynCom under either NC (white) or LP (grey). Col-0 roots under corresponding conditions were as control samples. MYC2 protein levels first were quantified by ACTIN then normalized to NC-BFO condition. Three independent biological replicates including four independent replicates of immunoblots (n = 32 samples). Letters indicate statistical significance (Kruskal-Wallis with Dunn’s *post hoc* test, α = 0.05). One immunoblots replicate was shown on the bottom. **(B)** Heatmaps showing the hierarchical relationship between samples from *jin1-8* pMYC2:MYC2-FLAG line (left) and *myc2-3* mutant (right) used for RNAseq. Three independent biological replicates (n = 48 samples). NC: normal light condition, LP: low PAR. -BFO: no microbes (i.e., germ-free), +BFO: with microbes. B: bacteria, F: fungi, O: oomycetes. The colour key on the top left side means sample distances (using *“dist’* function in “DESeq” R package to the transpose of the transformed count matrix to get sample-to-sample distances) corresponding to the colour range in the heatmap. **(C)** Bar plot showing the numbers of differentially regulated genes *myc2-3* mutant *vs. jin1-8* pMYC2:MYC2-FLAG line (|log2FC| ≥ 1, Empirical Bayes Statistics, FDR < 0.05). Red colour and blue colour filled bars represent significantly up- and down-regulated genes in the *myc2-3* mutant *vs. jin1-8* pMYC2:MYC2-FLAG line, respectively. The numbers highlighted inside the bars indicate the total number of up-or down-regulated genes in the *myc2-3* mutant *vs. jin1-8* pMYC2:MYC2-FLAG line. **(D)** Transcript profiling of 5,231 *A. thaliana* genes significantly regulated in root samples between *myc2-3* mutant and *jin1-8* pMYC2:MYC2-FLAG line (|log2FC| ≥ 1, Empirical Bayes Statistics, FDR < 0.05). Over-represented (white to red) and under-represented transcripts (white to blue) are shown across conditions as mean the differential gene expression between *myc2-3* and *jin1-8* pMYC2:MYC2-FLAG (log2 fold-change of counts per million (TMM-normalized voom-transformed data with limma package in R)). The gene set was split into eight major MYC2 differentially-expressed gene expression clusters in root, labelled MDER1 to MDER8. NC: normal light condition, LP: low PAR, -BFO: no microbes, +BFO: with microbes. Three independent biological replicates (n = 24 samples). **(E)** Overlap between the number of MDE genes identified in shoot (top) or root (below) and all MYC2 target genes identified by ChipSeq experiments in three independent studies (**Van Moerkercke et al., 2019; Wang et al., 2019; Zander et al., 2020**). **(F)** Gene Ontology (GO) term enrichment depicting the top 12 most significantly enriched GO terms (Hypergeometric test with Bonferroni correction, *P* < 0.05) detected in all clusters for root samples (left) and shoot samples (right). The size of point reflects the amount of gene numbers enriched in this GO term. The colour of point means the p value (Hypergeometric test with Bonferroni correction). R: response, Reg. Regulation, ET: ethylene, SA: salicylic acid, JA: jasmonic acid, SAR: systemic acquired resistance, CW: cell wall, CG: cell growth, P: process, Com: compound, Hex: hexakisphosphate, inv.: involved in. **(G)** Phytohormone levels of five-week-old *A. thaliana* Col-0 and *myc2-3* mutants grown under NC and LP, in the absence (-BFO) or presence of the BFO SynCom (+BFO), in the FlowPot system. Phytohormone levels were scaled to 0 – 1 individually. Three independent biological replicates are shown (in shoots: OPDA n = 87 samples, JA n = 87 samples, JA-Ile n = 88 samples, SA n = 87 samples; in roots: OPDA n = 56 samples, JA n = 56 samples, JA-Ile n = 56 samples, SA n = 55 samples). Statistical significance across conditions for each genotype is indicated with letters (Kruskal-Wallis with Dunn’s *post hoc* test, α = 0.05). NC: normal light condition, LP: low PAR. OPDA: 12-oxo-phytodienoic acid, JA: jasmonic acid, JA-Ile: Jasmonic Acid-Isoleucine, SA: free salicylic acid.

## SUPPLEMENTARY TABLES

**Table S1. Microbial culture collections used for microbiota reconstitution experiments**

**Table S2. PERMANOVA partitioning of microbial community assemblages**

**Table S3. RNA-Seq expression profiling of *A. thaliana* genes in root and shoot samples**

**Table S4. *A. thaliana* mutants used in this study**

**Table S5. PERMANOVA partitioning of microbial community assemblages across *A. thaliana* mutants**

**Table S6. RNA-Seq expression profiling of *A. thaliana* genes (*myc2-3 - jin1-8* pMYC2:MYC2-FLAG) in root and shoot samples**

